# Ancient Admixture into Africa from the ancestors of non-Africans

**DOI:** 10.1101/2020.06.01.127555

**Authors:** Christopher B. Cole, Sha Joe Zhu, Iain Mathieson, Kay Prüfer, Gerton Lunter

## Abstract

Genetic diversity across human populations has been shaped by demographic history, making it possible to infer past demographic events from extant genomes. However, demographic inference in the ancient past is difficult, particularly around the out-of-Africa event in the Late Middle Paleolithic, a period of profound importance to our species’ history. Here we present `SMCSMC`, a Bayesian method for inference of time-varying population sizes and directional migration rates under the coalescent-with-recombination model, to study ancient demographic events. We find evidence for substantial migration from the ancestors of present-day Eurasians into African groups between 40 and 70 thousand years ago, predating the divergence of Eastern and Western Eurasian lineages. This event accounts for previously unexplained genetic diversity in African populations, and supports the existence of novel population substructure in the Late Middle Paleolithic. Our results indicate that our species’ demographic history around the out-of-Africa event is more complex than previously appreciated.

## 1 Introduction

Methods to characterize demographic history from genetic data alone form a useful and independent complement to archaeological approaches [40], and many methods have been developed for this purpose [11, 14, 20–22, 30, 34, 43, 47, 52, 55, 56]. A particularly interesting time in human history is the end of the Middle Paleolithic (∼ 60 thousand years ago (kya)), which saw the divergence of the most deeply sampled lineages of human genetic variation, introgression from multiple archaic sources, and the expansion of anatomically modern humans Out of Africa (OoA). As archaeological evidence and ancient DNA from this period are scarce, inference of demography from present-day genetic data is potentially very informative, though technically challenging.

The phylogenetic trees over a set of samples as they change along the genome through recombination, collectively referred to as the ancestral recombination graph (ARG) [2, 21], record all information about the samples’ evolutionary history. This history itself is shaped by the population’s demography, a statistical relationship that is quantified by the coalescent-with-recombination (CwR) model [2]. The ARG is a complex data structure which is only weakly constrained by the observed genetic polymorphisms, making inference of demography difficult. By making approximations to the CwR, for instance by making an independent-sites assumption, efficient parametric inference of demography becomes possible [6, 18]. Methods including `PSMC` [11], `diCal` [30] and `SMC++` [31] allow non-parametric inference of demography under a closer approximation to the CwR, but one that does not include gene flow between populations. `MSMC` [22] introduced the cross-coalescent rate and `MSMC-IM` described how to interpret this rate in the context of a isolation-migration model to estimate a migration rate between populations[56]. However, these methods are not well suited for estimating directional migration rates.

Here we extend `SMCSMC` (Sequential Monte Carlo inference of the Sequentially Markovian Coalescent, [44]) to allow inference of directional migration. `SMCSMC` is a Bayesian method that uses a particle filter to explicitly sample from the posterior distribution of ARGs over multiple diploid samples under the full CwR model. Since particle filters operate by simulating latent variables (here the ARG) under the statistical model of interest, it becomes possible to handle complex demographic scenarios. We exploit this by extending the CwR model to include time-varying directional migration rates in a two-island demographic model. We use the posterior sample of ARGs including migration events to update the parameters of the demographic model, using either expectation-maximization or a variational Bayes procedure, and iterate these steps until convergence. We apply `SMCSMC` to estimate directional migration rates in whole genome sequencing data from the Simons Genome Diversity Panel (SGDP) [35] and the Human Genome Diversity Panel (HGDP) [48] to investigate population structure around the OoA event.

## 2 Results

### Substantial Migration from Eurasian to African Ancestors

We use `SMCSMC` to analyse pairs of individuals from the SGDP and simultaneously infer migration rates and effective population sizes (*N*_*e*_) under a two-island model with directional migration. Population sizes and migration rates are modeled as piece-wise constant across 32 exponentially spaced epochs from 133 to 133016 generations in the past, corresponding to 3.8 thousand to 3.8 million years ago (3.8kya–3.8Mya) using a generation time *g* = 29 years [5]. We find that the method infers high rates of migration from descendants of the OoA event (‘non-Africans’) to Africans, but not in the opposite direction, in the period 30–70kya corresponding to the Late Middle Paleolithic (Fig. 1). In populations from the Niger-Kordofanian and Nilo-Saharan language groups, comprising the majority of the population on the African continent, the peak inferred migration rate from Eurasian populations (2.5–3.0 × 10^−4^ and 3.5–4.0 × 10^−4^, in units of proportion of the target (ancestral African) population replaced per generation) most frequently falls in the epochs spanning 35–45kya, while peak migration rates in the opposite direction are substantially lower (0.5-1.0 × 10^−4^) and occur earlier, in the epochs spanning 55–70kya (Supplemental Fig. S1). Populations in the Afroasiatic language group show evidence of large amounts of directional migration in the Holocene (Supplemental Fig. S2), which is consistent with previous findings of relatively recent European introgression into these populations [32, 50].

**Figure 1:**
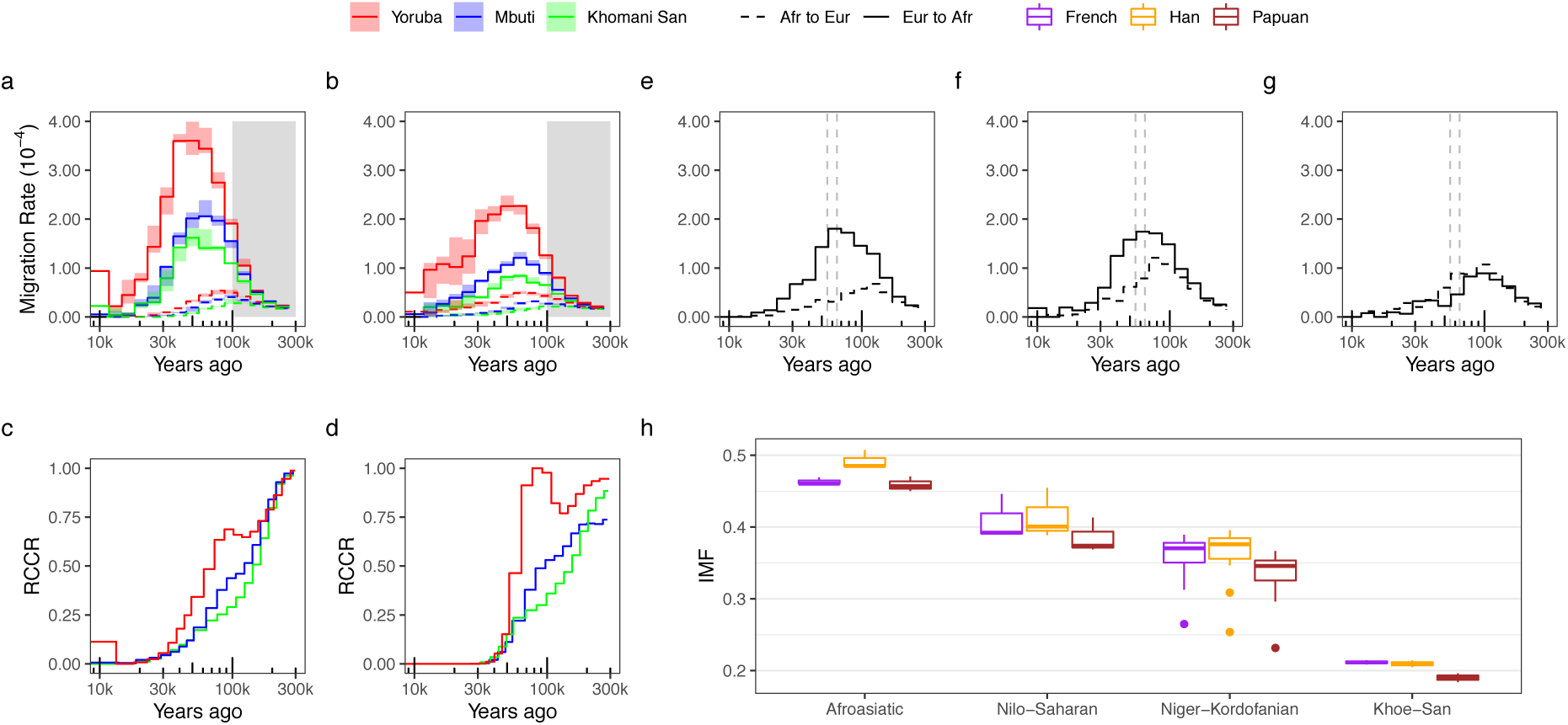
Migration rate inference. **a**. Inferred migration between an African individual and a Han Chinese individual in the Simons Genome Diversity Panel (SGDP) estimated using `SMCSMC`. Three replicates were performed, with the median estimate plotted and the range shaded. Solid lines show inferred migration from Eurasians to Africans (forward in time) while dotted lines show the reverse migration. The `SMCSMC` analysis used 10000 particles to estimate the posterior distribution of marginal trees, and 25 iterations of variational Bayesian inference to achieve converged parameter estimates. The shaded grey regions represents a time period where simulation shows `SMCSMC` has very little power to infer migration (Supplemental Section S5). The same analysis as in a. except using individuals from the physically phased subset of the Human Genome Diversity Panel (HGDP), showing similar differences between populations but systematically lower migration overall. Three replications were performed to estimate error and the standard deviation is shaded. The same `SMCSMC` settings were used as in a. **c**. Relative cross-coalescence rate (RCCR) estimated by `MSMC` in three different populations in the SGDP, supporting gene flow between Eurasians and Yorubans not shared by Mbuti or Khoe-San. 40 iterations were used to achieve parameter convergence. **d**. The same analysis as in c. but performed on individuals in the physically phased subset of the HGDP, similarly supporting shared gene flow between the Yoruban and Eurasians not shared by Mbuti or Khoe-San. **e, f** and **g**. Inferred migration rates from from data simulated under a two-island model with, from left to right, a backward Eurasia-to-Africa, a bidirectional, and a forward migration pulse lasting 10ky (dashed vertical lines) and replacing 40% of the recipient population(s) approximately 60kya. The migration rate from Africa to Eurasia is not well estimated by `SMCSMC` (see Figs. S8–S10 and Supplemental Section S5), but SMCSMC is well powered to infer migration from Eurasia to Africa in this period. **h**. Integrated total migration fraction (IMF) over the last 100 thousand years stratified by language phyla in the SGDP and comparison Eurasian population used to estimate migration. Afroasiatic (Mozabite, Saharawi, and Somali), Nilo-Saharan (Dinka, Luo, and Masai), Niger-Kordofanian (BantuHerero, BantuKenya, BantuTswana, Biaka, Esan, Gambian, Luhya, Mandenka, Mbuti, and Mende), and San (Khomani San and Ju hoan North) are grouped as in [50]. Similar levels of migration are inferred from French and Han Chinese to all language groups, with significantly less migration from Papuan groups (*p* ≤ 0.05, two-tailed paired t-test, Supplemental Table S1). Ourliers in the Niger-Kordofanian group are the Mbuti.

To assess the impact of errors introduced by statistical phasing, which was used to phase the SGDP data, we repeated the analyses above on a subset of physically phased individuals from the Human Genome Diversity Project (HGDP) [35] (Supplemental Section S3.2). This data set comprises individuals from four African (Yoruban, San, Mbuti, and Biaka) and nine non-African populations (Druze, Han, Karitiana, two Papuan populations, Pathan, Pima, Sardinian, and Yakut). `SMCSMC` results in the HGDP are qualitatively similar to those in the SGDP (Fig. 1b, Supplemental Fig. S1a). Inferred migration rates are, in general, lower in HGDP data than when using matched SGDP samples (Figs. 1a,b and S4, demographic inference in matched sampled in Supplemental Fig./ S3), but in all cases, the migration rates from Eurasia to Africa are substantially higher than in the opposite direction, consistent with the findings in the SGDP (Supplemental Fig. S4).

We asked whether `SMCSMC` has power to detect a large back-migration event in the Late Middle Paleolithic and distinguish it from other demographic scenarios. To answer this we used `SCRM` [29] to simulate a gigabase of sequence data under a two-island demographic model, with effective population sizes chosen to be comparable to typical African and Eurasian populations as inferred from real data. To this we added a 10ky pulse of forward, backward or bidirectional migration of varying strengths, with the midpoint of the migration pulse within the range 40 to 70kya. To quantify the inferred amount of migration we calculate the integrated migration fraction (IMF), defined as one minus the probability that a lineage in the destination (e.g. African) population traced backwards in time remains in that population across a given epoch according to the migration model (see Methods). For the simulations, we chose the most recent 100kya as epoch, and used scenarios with IMFs ranging from 0 to 0.593. For each simulation we report the inferred IMF in both the forward and backward direction (Fig. 2); full results are given in Supplemental Section S5. We find that `SMCSMC` has good power to detect backward migration pulses up to 60kya (median ratio of inferred and true IMF, 0.91), while power drops off at 70kya (IMF ratio 0.46). In the pure backward migration case, some forward migration is falsely inferred, but this is always substantially less than the inferred backward migration (median ratio inferred forward to true backward IMF, 0.37; true migration peak 60kya). However, in the case of true forward migration as well as bidirectional migration, roughly equal mixtures of forward and backward migration are inferred (Fig. 2). We conclude that in the epoch 40–70kya the forward and bidirectional scenarios are difficult to distinguish from each other, but both can be distinguished from backward migration, the only scenario resulting in substantially different inferred backward and forward migration.

**Figure 2:**
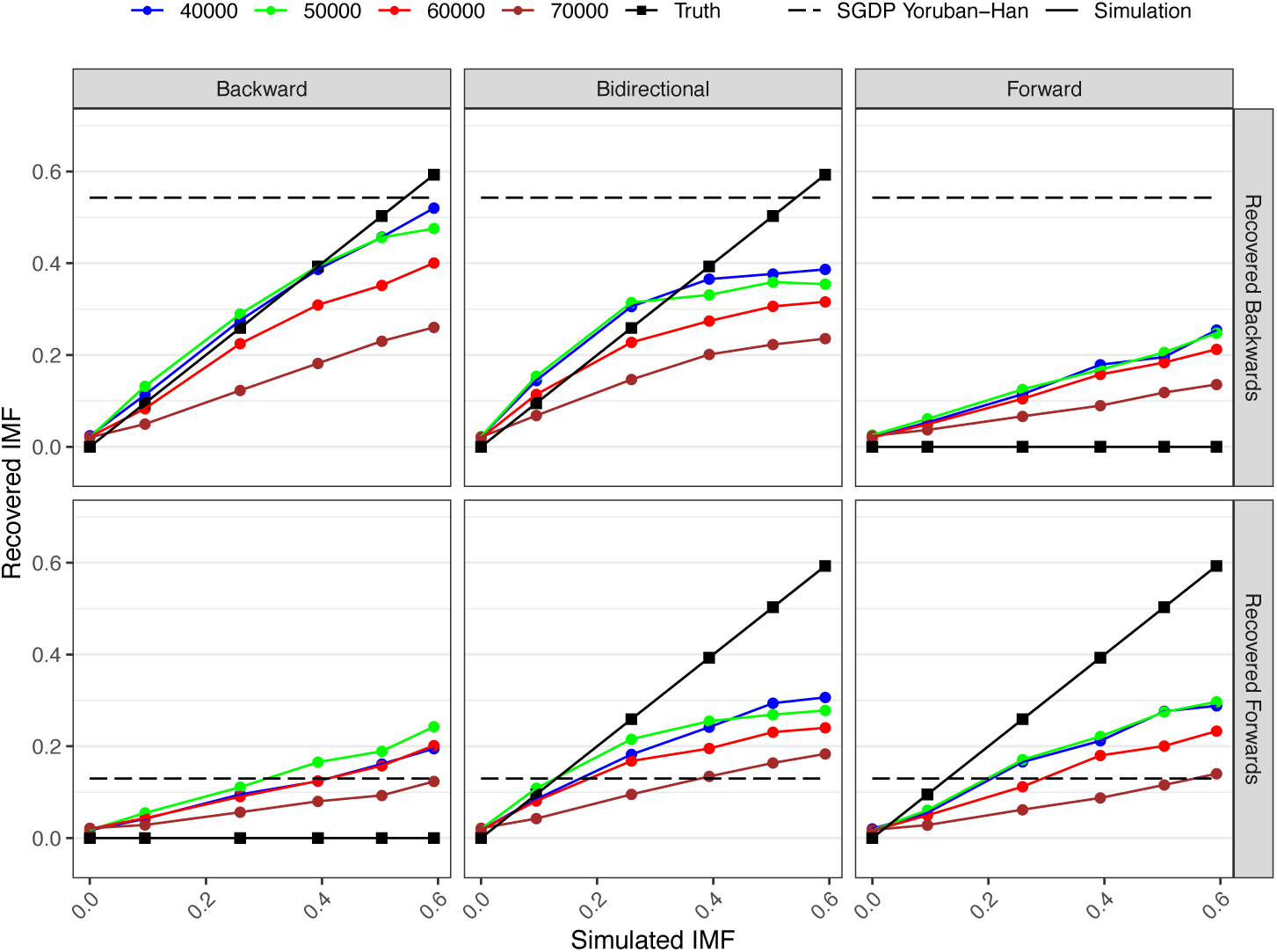
Simulation study. `SCRM` was used to simulate 1 gigabase sequence data for two diploid individuals under three different migration models. Migration was simulated backwards (from a Eurasian-like population to an African-like population), forward (the reverse), and symmetrically (equal migration in both directions). The amount of migration indicates the proportion of the sink population replaced by the source over a 10ky period centered at 40, 50, 60, or 70kya. The total IMF inferred by `SMCSMC` over the last 100ky is plotted and compared to the true simulated amount. For reference, the inferred IMF in either direction across 0-100kya for a Yoruban and Han individual is given in dashed lines. 5 iterations of variational Bayes and 5000 particles were used for inference. The effective population size model and additional details are given in Supplemental Section S5.

To validate the existence of the migration pulse, though not its direction, we next analyzed the same data using MSMC, which is widely used to estimate gene flow in the ancient past by estimating the relative cross-coalescent rate (RCCR) between two populations [22, 26, 28, 50]. We use the updated implementation MSMC2 recommended by the authors and first published in [34]. Each of the `SMCSMC` analyses are repeated using MSMC2 to estimate effective population size and RCCR (Supplemental Figs. S2, S4, S5). Consistent with previous analyses conducted with MSMC2, our estimates show high RCCR in the Late Middle Pleistocene in both the SGDP and the HGDP (Fig. 1c,d) [48, 50]. These observations confirm the existence of a substantial pulse of ancient gene flow between Eurasians (Han Chinese) and Africans.

### Migration Pre-dates East-West Eurasian Divergence

To assess whether the inferred back-migration shows variation across the descendants of the OoA event, we repeated the analyses using three representative non-African groups in the SGDP: Han Chinese, French European, and Papuans. Since simulations show that `SMCSMC` has little power to detect migration predating 70kya, and to exclude Holocene migration, the epoch we use to calculate real-data IMFs comprise the period of peak inferred migration up to the period of diminishing power (30–70kya); we use this epoch for all subsequent analyses. Inferred IMFs are not significantly different between Han Chinese and European populations in non-Afroasiatic populations (p=0.14, two-tailed paired t-test; Figs. 1h and S6, Table S1), consistent with migration occurring before the European-East Asian split approximately 40kya [20]. The contribution of this admixture event to extant African genetic variation is substantial; the estimated IMFs indicate that for individuals in the major African language groups, approximately a third of ancestral lineages trace their ancestry through the proto-Eurasian population (Niger-Kordofian group, 0.35 ± 0.04; Nilo-Saharan groups, 0.41 ± 0.03; Table S1). When we estimate these proportions using a Papuan sample to represent non-African descendants we find slightly but significantly smaller values compared to estimates using either the Han Chinese or European populations (mean difference of 0.029 ± 0.002, p=9.2 × 10^−15^, and 0.025 ± 0.004, p=2.3 × 10^−10^, paired t-tests, Supplemental Table S1, S2). Similarly, in the HGDP, inferred migration in both Papuan groups (Sepik and Highlands) was 0.025 ± 0.004 (p=1.4 × 10^−6^) lower than French and Han (Supplemental Table S3). We comment on this observation in the Discussion.

### Directional Migration Explains Excess Inferred African Genetic Diversity 100kya

Previous studies looking at effective population sizes (*N*_*e*_) in human ancestral populations have consistently reported inflated inferences in African populations approximately 100kya, often hypothesized to be due to unaccounted-for population substructure within Africa [11, 22]. We use `SMCSMC` to analyze African individuals paired with an individual from one of three non-African populations (Han Chinese, French European, and Papuans) and infer *N*_*e*_ for the African ancestral population under a two-island model with directional migration. Each analysis was repeated three times to assess the contribution of stochastic sampling to the inferences (Figs. 3, S5, per population *N*_*e*_ in Supplemental Fig. S7). `SMCSMC` infers substantially lower African *N*_*e*_ than `MSMC` in the period 80kya–300kya. In addition, while `MSMC` inferences show convergence of African and Eurasian ancestral *N*_*e*_ estimates only around 300kya, inferences from `SMCSMC` indicate convergence at 150kya (Fig. 3a), closer to the hypothesized time of the diversification of the ancestral lineages prior to the main out-of-Africa migration episode [34, 38]. The same analysis on physically phased samples from HGDP show that these results are not driven by errors due to statistical phasing (Fig. S4 and Supplemental Section S3.2). When we used `SMCSMC` to infer both African and European *N*_*e*_ under a single-population model without migration, *N*_*e*_ estimates were comparable to those from `MSMC` (Fig. 3b), indicating that the `SMCSMC` inferences are not driven by methodological biases particular to `SMCSMC`.

**Figure 3:**
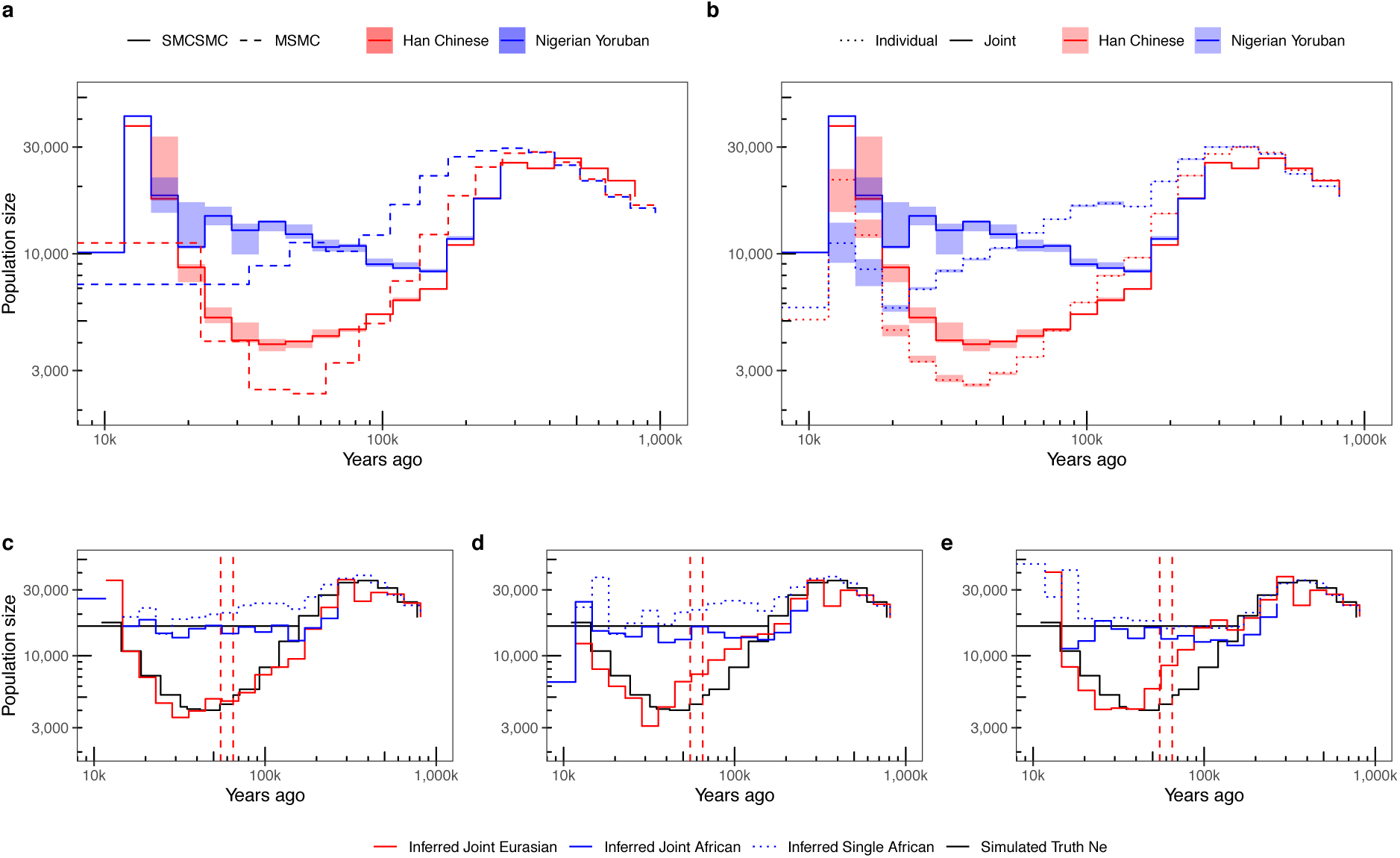
Effective population size inference. **a**. Analyzing a Nigerian Yoruban and a Han Chinese individual from the Simons Genome Diversity Panel jointly in a two-island model with directional migration using `SMCSMC` yields markedly lower *N*_*e*_ estimates and a more recent apparent split time, than when the same data are analyzed using `MSMC` with a model that does not explicitly include migration. Analyses for `SMCSMC` repeated three times; range of the estimates shaded. **b**. When each individual is analysed separately, using a model not including migration, *N*_*e*_ estimates from `SMCSMC` are similar to those of `MSMC2`. (Joint estimate from **a** included for comparison.) **c, d**, and **e**. Inferred Eurasian and African *N*_*e*_ from data simulated under a two-island model with, from left to right, a backward Eurasia-to-Africa, a bidirectional, and a forward migration pulse lasting 10ky (dashed vertical lines; same data as for Fig. 1**e**-**g**). Particularly for the backward migration case, inferred *N*_*e*_ under a two-island model tracks the true values (black) well, while inferred *N*_*e*_ under a single-population model are inflated around the split time. All `SMCSMC` analyses used 10000 particles and 25 variational Bayesian iterations; `MSMC` analyses used 40 iterations (Supplemental Section S5).

To more directly support the interpretation that the lower African *N*_*e*_ inferred by `SMCSMC` is due to appropriate modeling of directional migration, we again used coalescent simulation with `SCRM` to investigate various migration scenarios and their effects on inferred African *N*_*e*_. Using the simulation framework as above, we examine *N*_*e*_ estimates inferred under a two-island model with migration, and in addition *N*_*e*_ separately inferred for each of the two simulated populations under a single-population model (Supplemental Section S5). Focusing on single-population inferences, we found that for simulated African populations that had received substantial migration from the simulated Eurasian population either through backward or bidirectional migration, inferred *N*_*e*_ values indeed were substantially inflated compared to true values (Fig. 3c,d), while this effect was not seen when forward (African-to-Eurasian) migration was simulated (Fig. 3e). Similarly, single-population Eurasian *N*_*e*_ estimates were inflated in the presence of forward and bidirectional migration, but not backward migration (Supplemental Figs. S8–S10). In contrast, when using a model that includes migration, inferred African *N*_*e*_ do not show inflation in any of the three scenarios (Fig. 3c-e). We conclude that the inferences from `SMCSMC` and `MSMC` are compatible with substantial back-migration from ancestral Eurasians into Africans, but not substantial bidirectional or forward migration.

### Less Gene Flow to Central and South African Hunter-Gatherers

We infer substantial Eurasian back-migration into all African groups, however the inferred IMFs for individuals from Khoe-San populations are significantly lower than for any other group (difference with Niger-Kordofians, 0.14 ×0.02, *p* = 4.4 ×10^−14^; difference with Nilo-Saharans, 0.20 ± 0.03, *p* = 6.9 × 10^−9^, two-tailed t-test, Table S4). To further support this observation we used `MSMC` to estimate the relative cross-coalescent rate (RCCR) for several populations, and find evidence for gene flow between Yorubans and Eurasians that is not shared with the Khoe-San individuals in either the SGPD and the HGDP (Fig. 1c,d). These results are consistent across Eurasian donor populations (Fig. S3). The Khoe-San individuals are particular outliers, whose ancestors are inferred to have experienced approximately half the amount of admixture seen in Nilo-Saharan and Niger-Kordofanian groups (Fig. S6). In addition, we find that the Mbuti and Biaka, both Central African hunter-gatherer populations, show levels of Eurasian gene flow that are intermediate between levels observed in the Khoe-San and Yorubans (Fig. 1a,b, Supplemental Table S1). This is mirrored by inferred IMFs for Central African Hunter Gatherers, which are significantly lower than other Niger-Kordofanian groups (difference − 0.08 ±0.03, *p* = 1.2 × 10^−3^, Table S4), possibly reflecting the proposed early split times of the Mbuti and Biaka from the remainder of ancestral African populations between 60 and 200kya [41, 53].

### No Evidence for Excess Neanderthal Ancestry

Previous studies have proposed that a backflow from Eurasia may have brought Neanderthal ancestry into African populations [57]. To assess whether the proposed Late Middle Paleolithic back migration might have introduced Neanderthal material, we analyzed a Yoruban and a French individual using `SMCSMC` to draw a sample from the posterior distribution of ARGs, isolated the marginal trees containing an inferred back-migration event in the epoch 30–70kya, and reported the inferred admixture tracts (“segments”, Supplementary Section S4). To assess whether the identified segments are plausible, we confirmed that their length distribution is consistent with IMF and timing of the migration inferred by `SMCSMC` (Supplemental Section S4.1, S11), and, as expected, we found that these African segments with putative Eurasian ancestry tend to be more closely related to a Eurasian sample than another representative of the same African population (Table S5, Supplemental Fig. S12, S13) in a global dataset of modern and ancient individuals compiled by the Reich group (see URLs). Within these African segments that are likely enriched for material with Eurasian ancestry, we then used *D* statistics [13] to identify enrichment for Neanderthal material compared to an African background. We find no evidence for gene flow with a Vindija Neanderthal on the Mbuti baseline, or when compared to a different Yoruban (Table S6, S7). We additionally find no evidence for increased affinity to the Vindija Neanderthal when compared to the Altai, as would be expected if the material were descended from admixing Eurasians (Table S8). However, we find that restricted to the identified segments, *D* statistics have power to detect evidence for the known admixture from Vindija into a French individual (Fig. 4), suggesting that lack of power does not explain the lack of evidence we find for Neanderthal admixture into Africans. In addition, we find no differences in affinity to Neanderthals or Denisovans between the variants which fall in segments and the whole genome (Fig. 4d). Taken together, this suggests that Eurasian-derived segments of the African genomes are not enriched with Neanderthal material.

**Figure 4:**
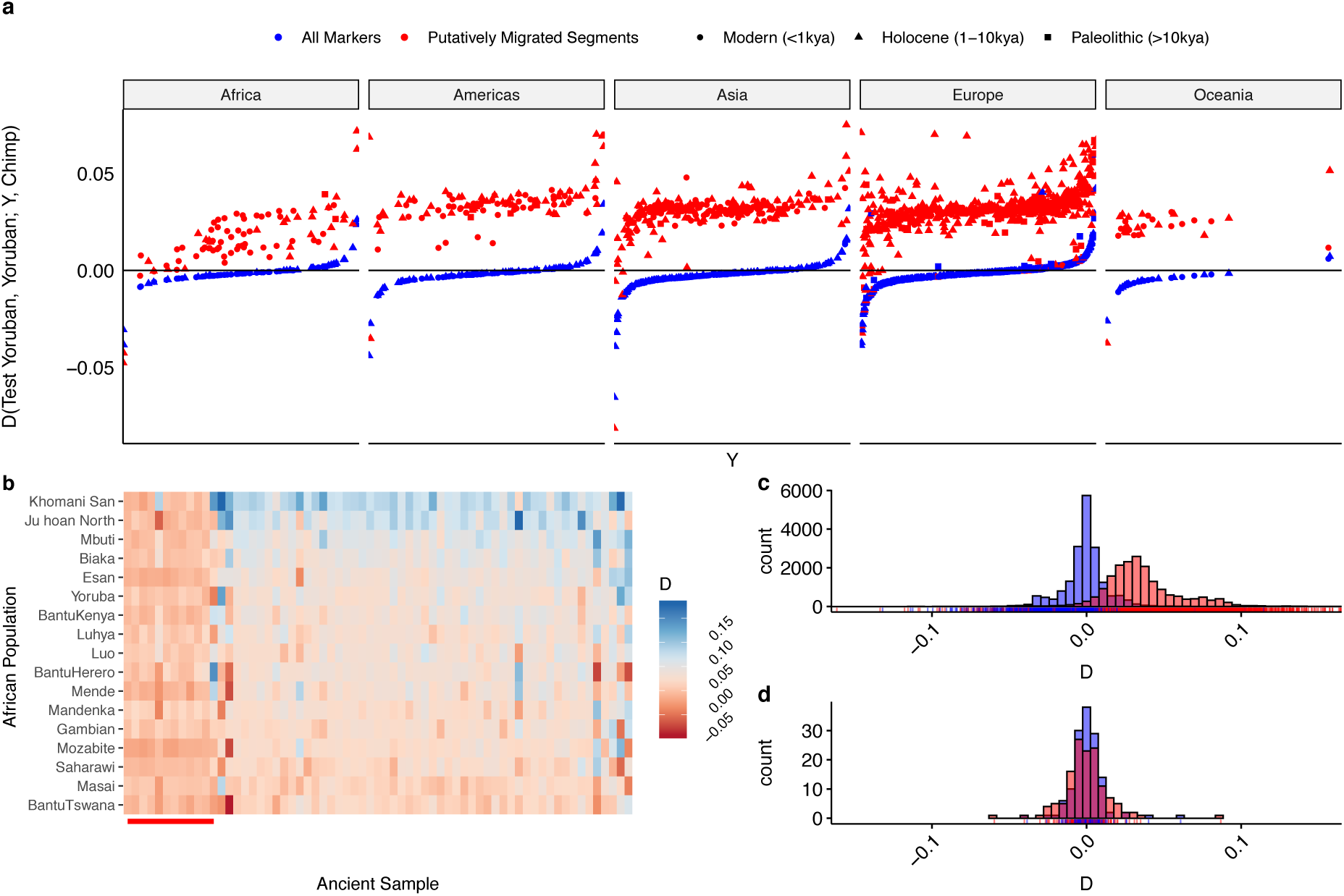
African introgressed segments are more similar to Eurasians but show no Neanderthal or Denisovan enrichment. **a**. *D*(Test Yoruban, Comparison Yoruban; Test population, Chimpanzee) calculated for all populations in the Reich Human Origins dataset (see URLs). *D* statistics in the putatively migrated segments are higher across the board in 3589 ancient, 6472 present day individuals. **b**. The same *D* statistic but computed for all African populations and individuals sampled from the Paleolithic. Neanderthal and Denisovan samples (marked with red bar) show low affinity to a Yoruban in putatively migrated segments. Histogram of *D* statistics computed in a. showing clear inflation of statistics calculated in segments (red) versus all markers (blue). **d**. Subset of individuals from a. involving Neanderthal (*n* = 6), Denisovan (*n* = 1), and a unique mixture individual (*n* = 1) with statistics calculated in segments (red) and all markers (blue) for all *n* = 17 African individuals indicating no difference in this population.

**Figure 5:**
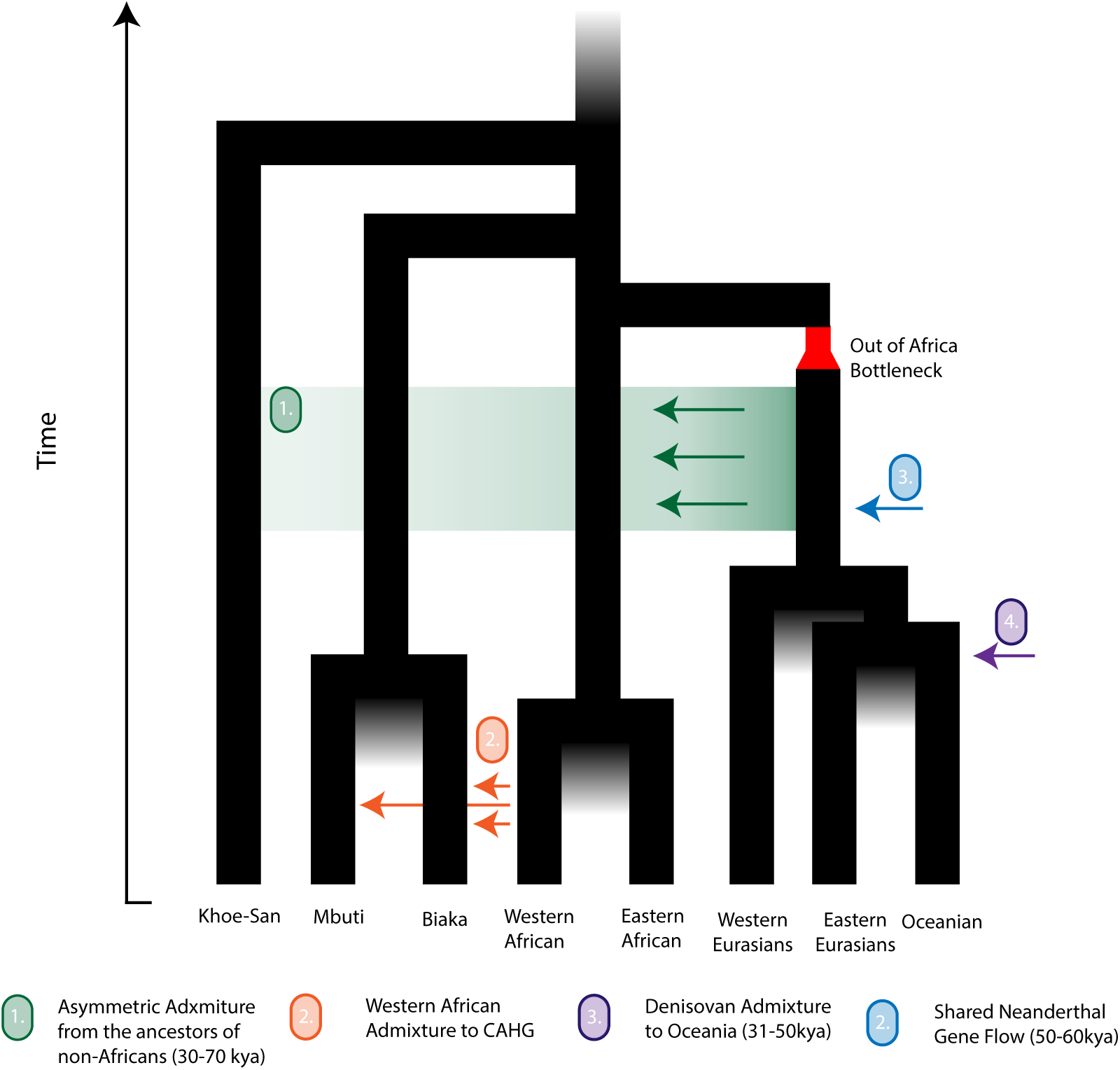
Proposed model for the diversification of modern human lineages. The proposed admixture from ancestors of present-day non-Africans is indicated in green. Several elements of this model remain unresolved, including the timing of the proposed admixture into Central and Southern Hunter Gatherer populations and these population’s branching order, gene flow from an unsampled archaic African human lineage, and the different episodes of Neanderthal and Denisovan admixture. Date ranges for the diversification of Central and Southern African Hunter Gatherers [35], Out of Africa migration [53], Eastern / Western Eurasian diversification [22], Eastern / Western African diversification [50], Oceanian / Eastern Eurasian diversification [39], the split of the Mbuti from Biaka [35], Neanderthal gene flow [15], and Denisovan Gene flow [34] may be found in references given.

## 3 Discussion

We have developed an approach for estimating demographic parameters and ARGs from whole genome sequence data, which can handle inference in complex demographic models, and implemented this in the software program `SMCSMC` [44]. We used `SMCSMC` to investigate ancient migration rates and population substructure, and found evidence for a substantial admixture from ancestors of present-day Eurasian populations into African populations in the Late Middle Paleolithic.

Our analysis suggests that a population ancestral to present-day Eurasians contributed as much as a third of the genetic material in many modern African populations. We find no difference in inferred admixture proportions when using French Europeans or Han Chinese as extant representatives of the donor population, indicating that the admixing population must have split from the out-of-Africa population before the East/West Eurasian divergence, implying a lower bound on the timing of the admixture of approximately 40kya [20]. It appears that our results suggest that the migrating population was more similar to present-day French and Chinese populations than to Papuans. However, up to 5% of the genomes of some present-day Papuans have been suggested to derive from archaic introgressions [37], and these contributions will have reduced the inferred levels of admixture into Africans when using Papuans as a representative of the Eurasian ancestors. The alternative explanation, of an earlier divergence of Papuans and Eurasian ancestors, is possible but contested; in light of documented Eurasian admixture into Oceania, the effects of this early isolation are likely to be small relative to the large confounding effects of Denisovan admixture [34, 40].

The proposed period of admixture has biased previous inferences of the African population sizes. We show that including directional migration into the model resolves previously unexplained high inferred *N*_*e*_ in the period 80 to 300kya. It is well known that effective population size estimates are biased in the presence of population substructure and migration [11, 43]. We use simulations to show that the proposed admixture event indeed causes an increase in estimated *N*_*e*_ in analyses that do not explicitly model migration. Correctly modeling of directional migration recovers the correct *N*_*e*_, and allows us to infer a more recent split time between the two populations than indicated by previous analyses, although we did not attempt to formally estimate this time of divergence.

We found that not all populations in Africa have been equally affected by the proposed migration event. While the ancestors of Niger-Kordofanian and Nilo-Saharan populations show evidence of similar levels of Eurasian admixture, the ancestors of Central African and South African hunter-gatherer populations show markedly lower levels. The date of genetic diversification of both the Central Hunter Gatherers and Khoe-San (SAHG) is contested [53], but a date of 100kya has been proposed [17], providing a putative upper bound on the main admixture event. Our simulations indicate that `SMCSMC` has little power to detect the impact of migration events occurring more than 70kya, providing an additional upper bound on the time of the migration episode, or the fraction of it that left a sufficiently distinct imprint on extant genetic material.

Compared to the remainder of the Niger-Kordofanians and Nilo-Saharans on the one hand, and the SAHG populations on the other, the Mbuti and Biaka show intermediate levels of admixture. Of these populations, the Biaka show slightly higher levels of admixture than the Mbuti, which is likely due to the well-documented admixture from Western African groups not shared with the Mbuti [10]. The lower levels of admixture in Mbuti and Biaka compared to Niger-Kordofian and Nilo-Saharan populations imply at least partial diversification of the former at the time of the migration, placing an upper bound on the timing. However, dating the diversification of these groups is difficult. Recent estimates using *f* statistics place the split concurrent with the San in a large-scale early expansion 200-250kya [53], while older data consistently report an earlier split time between 50 and 90 kya [45]. Further clarity on the early structure and diversification of hunter-gatherer populations are necessary to interpret their interactions with Eurasian migrants. The Afroasiatic populations on the other hand show high levels of admixture, which also appears to be of much more recent origin, and it appears likely that this is the result of extensive admixture from Eurasian populations during the Holocene [32, 50].

It has previously been suggested that Eurasian back-migration may be responsible for Neanderthal material in Africans [57]; however, we find no evidence for enrichment of Neanderthal-like material in putatively Eurasian-derived genomic segments in Africans, indicating that Neaderthal introgression into Eurasians occurred after the African introgression event we study here, or that further population structure in the Eurasian ancestral population precluded substantial transmission of Neanderthal material into Africa.

Our findings are consistent with several other published observations. Migration rate estimates using `MSMC-IM` revealed high levels of admixture at times comparable to our results [56]. The coalescent intensity function additionally shows similar histories between sub-Saharan African and Eurasian groups with high coalescent intensity in epochs consistent with our inference and those of `MSMC-IM`, supporting both an early split between the groups and a substantial replacement of genetic material more recently than 100kya [47]. Evidence has been mounting for multiple migrations into the Eurasian continent, possibly mediated by climatic drivers [36, 38]. Eurasian backflow during the Holocene has been well established [24, 25], but earlier migrations have also been proposed before based on observations of the spatial distribution of Y chromosome and mitochondrial haplogroups [1, 3, 4, 8, 33, 42, 51]. At the same time, evidence has been mounting for extreme heterogeneity in the history of sub-Saharan Africans, with several unsampled population theorised to have contributed at various points in the past [49, 53, 55]. In light of these recent studies, the observations in this paper add to a growing body of evidence for complex population structure and migration surrounding the Out of Africa event leading to a substantial replacement of the African population in the Late Middle Paleolithic.

## 4 Methods

### A Particle Filter for Demographic Inference

Details of the Sequential Monte Carlo for the Sequentially Markovian Coalescent (`SMCSMC`) algorithm have been previously published [44] (see the URLs for an implementation). Briefly, `SMCSMC` builds an approximation of the posterior distribution of genealogical trees along the genome using a particle filter, a method also known as sequential Monte Carlo. It does so by simulating a number of sequences of genealogical trees (particles) under a fixed set of demographic parameters *θ*, using the sequential coalescent sampler `SCRM` [29]. Simulated recombination events may change the local trees along the sequence. Particles are then weighted according to their conditional likelihood given observed polymorphisms. To avoid sample depletion, the set of particles is regularly resampled, which tends to remove and duplicate particles with low and high weight respectively. To further increase the efficiency of the procedure, the resampling procedure targets not the partial posterior distribution that includes polymorhpisms up to the current location, but also includes a “lookahead likelihood” term that approximates a particle’s likelihood’s dependence on subsequent polymorphisms, while ensuring that the estimate of the posterior tree distribution remains asymptotically exact. From a sample of trees from the posterior distribution, Variational Bayes (VB) or Stochastic Expectation Maximization (SEM) is used to update the estimates of demographic parameters *θ*. This is repeated over a given number of iterations, or until *θ* have converged.

To add the ability to infer time-varying migration rates, we exploit the capabilities of `SCRM` to simulate ARGs under complex demographic scenarios, and collect sufficient statistics (migration opportunity, and number, time and direction of simulated migration events) for each particle.

We use `SMCSMC` to infer effective population sizes and migration matrices in pairs of unrelated individuals from the phased release of the Simons Global Diversity Panel. We set a uniform recombination rate of 3 ×10^−9^ and a neutral mutation rate of 1.25 ×10^−8^, both in units of events per nucleotide per generation; previous results indicate that modeling recombination hotspots minimally affects results [11]. To reduce the number of iterations to convergence, we initialise the particle filter with an approximation of human demographic history (Supplemental Fig. S14). We seed the model with an initial constant symmetric migration rate of 0.0092 (*M*_*i,j*_; proportion per generation of the sink population replaced by migrants from the source backwards in time). We arrive at this value through simulation (Supplemental Section S5, Supplemental Figs. S15, S16).

### Multiply Sequential Markovian Coalescent

We use `MSMC2` to estimate the effective population size of pairs of African and Eurasian individuals using default configurations and scripts provided in `msmc-tools` (see URLs) [22, 56]. We use a fixed recombination rate in line with our `SMCSMC` analysis and skip ambiguously phased sites. Twenty iterations are performed by default. We additionally compute the relative cross-coalescent rate to examine relative gene flow by transforming the coalescent rates generated by `MSMC2` as indicated in the software documentation.

### Coalescent Simulation

Coalescent simulations were performed under the sequential coalescent with recombination model (`SCRM`) [29]. Full details of the simulation procedure are detailed in Supplemental Section S5. 1 gigabase (Gb) of sequence was simulated. In addition to branches in local genealogical trees, `SCRM` retains non-local branches in the ancestral recombination graph (ARG) within a user-specified sliding window. In the limit of a chromosome-sized windows `SCRM` is equivalent to the coalescent with recombination, while for a zero-length window it is equivalent to the sequentially Markovian coalescent (SMC’) [6, 7]; we use a 100kb sliding window to approximate the CwR and improve accuracy over SMC’ while retaining tractable inference.

We modelled migration as a 10ky pulse of constant migration rate resulting in an integrated migration fraction (IMF) of 0 to 0.593. The migration pulse was centered at various times between 40 and 70 kya. Due to the amount of compute required, we then used `SMCSMC` to infer the demographic parameters using a reduced set of 5000 particles and 5 iterations of the VB procedure. To aid convergence, we started inference at a reasonable approximation of human demographic history (see Supplemental Section S5.1, Supplemental Fig. S17). We modelled *N*_*e*_ and migration rates as piecewise continuous functions and set 32 exponentially spaced epochs from 133 to 133016 generations in the past. To convert evolutionary rates to years we set a generation time of 29 years [5]. For computational efficiency, individual genomes were split into 120 chunks and processed in parallel, with sufficient statistics collected and processed together in the VB steps.

### Isolating Anciently Admixed Segments

We sampled genealogical trees with migration events from the posterior distribution estimated by the particle filter under the final, converged, demographic parameters. We scan along the sequence and identified marginal trees with migration events from the source (Eurasian) population to the sink (African) population (forward in time) within the desired time period along with the beginning and end position of that tree in the genome sequence. In this process, we ignore recombination event that alter a tree in such a way that the migration event is retained.

### Sequence Data and Preparation

We downloaded whole genome sequence (WGS) data from the phased release of the Simons Genome Diversity Panel and converted it to `.seg` file format using scripts provided (See URLs). We apply two masks to the data. First, we mask the data with the strict accessibility mask provided by the 1000 genomes project (see URLs). Second, we mask any sites absent chimpanzee ancestry, to address a known variant issue in the data that resulted in artificially long runs of homozygosity [56]. We develop a `Snakemake` [12] pipeline for efficiently analysing sequence data with both `SMCSMC` and `MSMC2`. We assume a mutation rate of 1.25 × 10^−8^ and a recombination rate of 3 × 10^−9^ (events per nucleotide per generation), in line with recent literature [16, 23]. The number of particles, and the number of VB iterations, are set per analyses, and are reported in figure captions. Unless otherwise noted, the names of individuals used in this paper are the first in their population (e.g. an individual named Yoruban is `S Yoruba-1` in the SGDP nomenclature); a complete list of sample identifiers is provided in Supplemental Table S9.

### Formal Statistics

Patterson’s formal statistics were calculated with `ADMIXTOOLS` [13] and the `admixr` package [54] in `R`. We converted the above sequence data to Eigenstrat format with `vcf2eigenstrat` formerly distributed with `admixr`. We merged SGDP and archaic Eigenstrat datasets with `convertf` and `mergeit` implemented in `ADMIXTOOLS`.

### Integrated Migration Fraction

The IMF, the total fraction of a particular population *A* replaced during a particular time period from *T*_0_ to *T*_1_ generations in the past is found as follows. Let *ρ* (*t*) be the instantaneous rate of migration out of *A* per unit of time in the backward direction (i.e. into *A* forwards in time), and *F* (*t*) the fraction not migrated in the epoch [*T*_0_, *t*], then 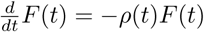 with solution 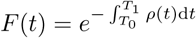, so that the IMF is given by 1 − *F* (*T*_1_). The integral is calculated as a finite sum since *ρ* is piecewise constant.

## 4.1 Acknowledgements

CBC acknowledges grant support from a Wellcome Trust studentship 215098/Z/18/Z. KP acknowledges support from the Max Planck Society. IM was supported by a Research Fellowship from the Alfred P. Sloan foundation (FG-2018-10647) and a New Investigator Research Grant (KA2018-98559) from the Charles E. Kaufman Foundation. Computation used the BMRC facility, a joint development between the Wellcome Centre for Human Genetics and the Big Data Institute supported by the NIHR Oxford BRC. The views expressed are those of the author(s) and not necessarily those of the NHS, the NIHR or the Department of Health. Financial support was provided by the Wellcome Trust Core Award (grants 090532/Z/09/Z and 203141/Z/16/Z), and the Strategic Alliance Funding: MRC Weatherall Institute of Molecular Medicine (WIMM) grant MC UU 12025.

## URLs

Simons Genome Diversity Panel phased release, https://sharehost.hms.harvard.edu/genetics/reich_lab/sgdp/phased_data/.

Human Genome Diversity Panel, ftp://ngs.sanger.ac.uk/production/hgdp/hgdp_wgs.20190516/.

Ancient DNA, http://cdna.eva.mpg.deandertal/.

Strict 1000 Genomes accessibility mask, ftp://ftp.1000genomes.ebi.ac.uk/vol1/ftp/release/20130502/supporting/accessible_genome_masks/.

`SMCSMC`, https://github.com/luntergroup/smcsmc.

`MSMC2`, https://github.com/stschiff/msmc2.

`msmc-tools`, https://github.com/stschiff/msmc-tools.

`vcf2eigenstrat`, https://github.com/bodkan/vcf2eigenstrat.

`ADMIXTOOLS`, https://github.com/DReichLab/AdmixTools.

`admixr`, https://github.com/bodkan/admixr.

## References

1. Altheide, T. K. & Hammer, M. F. Evidence for a possible Asian origin of YAP + Y chromosomes. American journal of human genetics 61, 462–6 (1997).

2. Griffiths, R. C. & Marjoram, P. in (1997).

3. Hammer, M. F. et al. Out of Africa and back again: nested cladistic analysis of human Y chromosome variation. Molecular Biology and Evolution 15, 427–441 (1998).

4. Cruciani, F. et al. A Back Migration from Asia to Sub-Saharan Africa Is Supported by High-Resolution Analysis of Human Y-Chromosome Haplotypes. The American Journal of Human Genetics 70, 1197–1214 (2002).

5. Fenner, J. N. Cross-cultural estimation of the human generation interval for use in genetics-based population divergence studies. American Journal of Physical Anthropology 128, 415–423 (2005).

6. McVean, G. A. & Cardin, N. J. Approximating the coalescent with recombination. Philosophical Transactions of the Royal Society B: Biological Sciences (2005).

7. Marjoram, P. & Wall, J. D. Fast “coalescent” simulation. BMC Genetics 7, 16 (2006).

8. Chandrasekar, A. et al. YAP insertion signature in South Asia. Annals of Human Biology 34, 582–586 (2007).

9. Dumont, B. L. & Payseur, B. A. Evolution of the genomic rate of recombination in mammals. Evolution 62, 276–294 (2008).

10. Batini, C. et al. Insights into the demographic history of African pygmies from complete mitochondrial genomes. Molecular Biology and Evolution 28, 1099–1110 (2011).

11. Li, H. & Durbin, R. Inference of human population history from individual whole-genome sequences. Nature 475, 493–496. arXiv: 1011.1669v3 (2011).

12. Köster, J. & Rahmann, S. Snakemake-a scalable bioinformatics workflow engine. Bioinformatics (2012).

13. Patterson, N. et al. Ancient Admixture in Human History. 192, 1065–1093 (2012).

14. Pickrell, J. K. et al. The genetic prehistory of southern Africa. Nature Communications 3, 1–6. 1207.5552 (2012).

15. Sankararaman, S., Patterson, N., Li, H., Pääbo, S. & Reich, D. The Date of Interbreeding between Neandertals and Modern Humans. PLoS Genetics (2012).

16. Scally, A. & Durbin, R. Revising the human mutation rate: Implications for understand ing human evolution. Nature Reviews Genetics 13, 745–753. issn: 14710064 (2012).

17. Schlebusch, C. M. et al. Genomic Variation in Seven Khoe-San. 1187, 374–379 (2012).

18. Excoffier, L., Dupanloup, I., Huerta-Sánchez, E., Sousa, V. C. & Foll, M. Robust Demographic Inference from Genomic and SNP Data. PLoS Genetics (2013).

19. Liang, M. & Nielsen, R. The Lengths of Admixture Tracts. Genetics 197, 953–967 (2014).

20. Mathieson, I. & McVean, G. Demography and the Age of Rare Variants. PLoS Genetics 10. arXiv: 1401.4181 (2014).

21. Rasmussen, M. D., Hubisz, M. J., Gronau, I. & Siepel, A. Genome-Wide Inference of Ancestral Recombination Graphs. PLoS Genetics (2014).

22. Schiffels, S. & Durbin, R. Inferring human population size and separation history from multiple genome sequences. Nature Genetics 46, 919–925. arXiv: 005348 [10.1101] (2014).

23. Schiffels, S. & Durbin, R. Inferring human population size and separation history from multiple genome sequences. Nature Genetics 46, 919–925. arXiv: 005348 [10.1101] (2014).

24. Gallego Llorente, M. & Manica, A. Ancient Ethiopian genome reveals extensive Eurasian admixture in Eastern Africa. Science 350, 820–825 (2015).

25. López, S., van Dorp, L. & Hellenthal, G. Human Dispersal Out of Africa: A Lasting Debate. Evolutionary bioinformatics online 11, 57–68 (2015).

26. Pagani, L. et al. Tracing the Route of Modern Humans out of Africa by Using 225 Human Genome Sequences from Ethiopians and Egyptians. American Journal of Human Genetics 96, 986–991 (2015).

27. Racimo, F., Sankararaman, S., Nielsen, R. & Huerta-Sánchez, E. Evidence for archaic adaptive introgression in humans. Nature Reviews Genetics 16, 359 (2015).

28. Raghavan, M. et al. Genomic evidence for the Pleistocene and recent population history of Native Americans. Science 349 (2015).

29. Staab, P. R., Zhu, S., Metzler, D. & Lunter, G. SCRM: efficiently simulating long sequences using the approximated coalescent with recombination. Bioinformatics 31, 1680–1682 (2015).

30. Steinrücken, M., Kamm, J. A. & Song, Y. S. Inference of complex population histories using whole-genome sequences from multiple populations. bioRxiv, 026591 (2015).

31. Terhorst, J. & Song, Y. S. Fundamental limits on the accuracy of demographic inference based on the sample frequency spectrum. Proceedings of the National Academy of Sciences 112, 7677–7682 (2015).

32. Busby, G. B. et al. Admixture into and within sub-Saharan Africa. eLife (2016).

33. Hervella, M. et al. The mitogenome of a 35,000-year-old Homo sapiens from Europe supports a Palaeolithic back-migration to Africa. Scientific Reports 6, 25501 (2016).

34. Malaspinas, A. S. et al. A genomic history of Aboriginal Australia. Nature 538, 207–214. arXiv: NIHMS150003 (2016).

35. Mallick, S. et al. The Simons Genome Diversity Project: 300 genomes from 142 diverse populations. Nature 538, 201–206 (2016).

36. Pagani, L. et al. Genomic analyses inform on migration events during the peopling of Eurasia. Nature 538, 238–242 (2016).

37. Sankararaman, S., Mallick, S., Patterson, N. & Reich, D. The Combined Landscape of Denisovan and Neanderthal Ancestry in Present-Day Humans. Current Biology 26, 1241–1247. issn: 09609822. http://dx.doi.org/10.1016/j.cub.2016.03.037 (2016).

38. Timmermann, A. & Friedrich, T. Late Pleistocene climate drivers of early human migration. Nature 538, 92–95 (2016).

39. Lipson, M., Reich, D. & Townsend, J. P. A working model of the deep relationships of diverse modern human genetic lineages outside of Africa. Molecular Biology and Evolution 34, 889–902. issn: 15371719 (2017).

40. Nielsen, R. et al. Tracing the peopling of the world through genomics 2017.

41. Patin, E. et al. Dispersals and genetic adaptation of Bantu-speaking populations in Africa and North America. Science 356, 543–546 (2017).

42. Cabrera, V. M., Marrero, P., Abu-Amero, K. K. & Larruga, J. M. Carriers of mitochondrial DNA macrohaplogroup L3 basal lineages migrated back to Africa from Asia around 70,000 years ago. BMC Evolutionary Biology 18, 98 (2018).

43. Chikhi, L. et al. The IICR (inverse instantaneous coalescence rate) as a summary of genomic diversity: insights into demographic inference and model choice. Heredity 120, 13–24 (2018).

44. Henderson, D., Zhu, S. (& Lunter, G. Demographic inference using particle filters for continuous Markov jump processes. bioRxiv, 382218 (2018).

45. Patin, E. & Quintana-Murci, L. The demographic and adaptive history of central African hunter-gatherers and farmers. Current Opinion in Genetics and Development 53, 90–97 (2018).

46. Zhou, Y., Tian, X., Browning, B. L. & Browning, S. R. POPdemog: visualizing population demographic history from simulation scripts. Bioinformatics (Oxford, England) issn: 13674811 (2018).

47. Albers, P. K. & McVean, G. Dating genomic variants and shared ancestry in population-scale sequencing data. bioRxiv, 416610 (2019).

48. Bergström, A. et al. Insights into human genetic variation and population history from 929 diverse genomes. bioRxiv, 674986 (2019).

49. Durvasula, A. & Sankararaman, S. Recovering signals of ghost archaic introgression in African populations. bioRxiv, 285734 (2019).

50. Fan, S. et al. African evolutionary history inferred from whole genome sequence data of 44 indigenous African populations. Genome Biology (2019).

51. Haber, M. et al. A Rare Deep-Rooting D0 African Y-Chromosomal Haplogroup and Its Implications for the Expansion of Modern Humans out of Africa. Genetics, genetics.302368.2019 (2019).

52. Kelleher, J. et al. Inferring whole-genome histories in large population datasets. Nature Genetics 51, 1330–1338 (2019).

53. Lipson, M. et al. Ancient West African foragers in the context of African population history (2019).

54. Petr, M., Vernot, B. & Kelso, J. admixr — R package for reproducible analyses using ADMIXTOOLS. Bioinformatics, 1–2 (2019).

55. Speidel, L., Forest, M., Shi, S. & Myers, S. R. A method for genome-wide genealogy estimation for thousands of samples. Nature Genetics 51, 1321–1329 (2019).

56. Wang, K., Mathieson, I., O’connell, J. & Schiffels, S. Tracking human population structure through time from whole genome sequences. bioRxiv, 1–21 (2019).

57. Chen, L. et al. Identifying and Interpreting Apparent Neanderthal Ancestry in African Individuals Article Identifying and Interpreting Apparent Neanderthal Ancestry in African Individuals. Cell, 1–11 (2020).

