## Supplemental Material for "Ancient Admixture into Africa from the ancestors of non-Africans"

Supplementary Material

**S5 Simulation procedure** **45**

### S1 Supplemental Figures

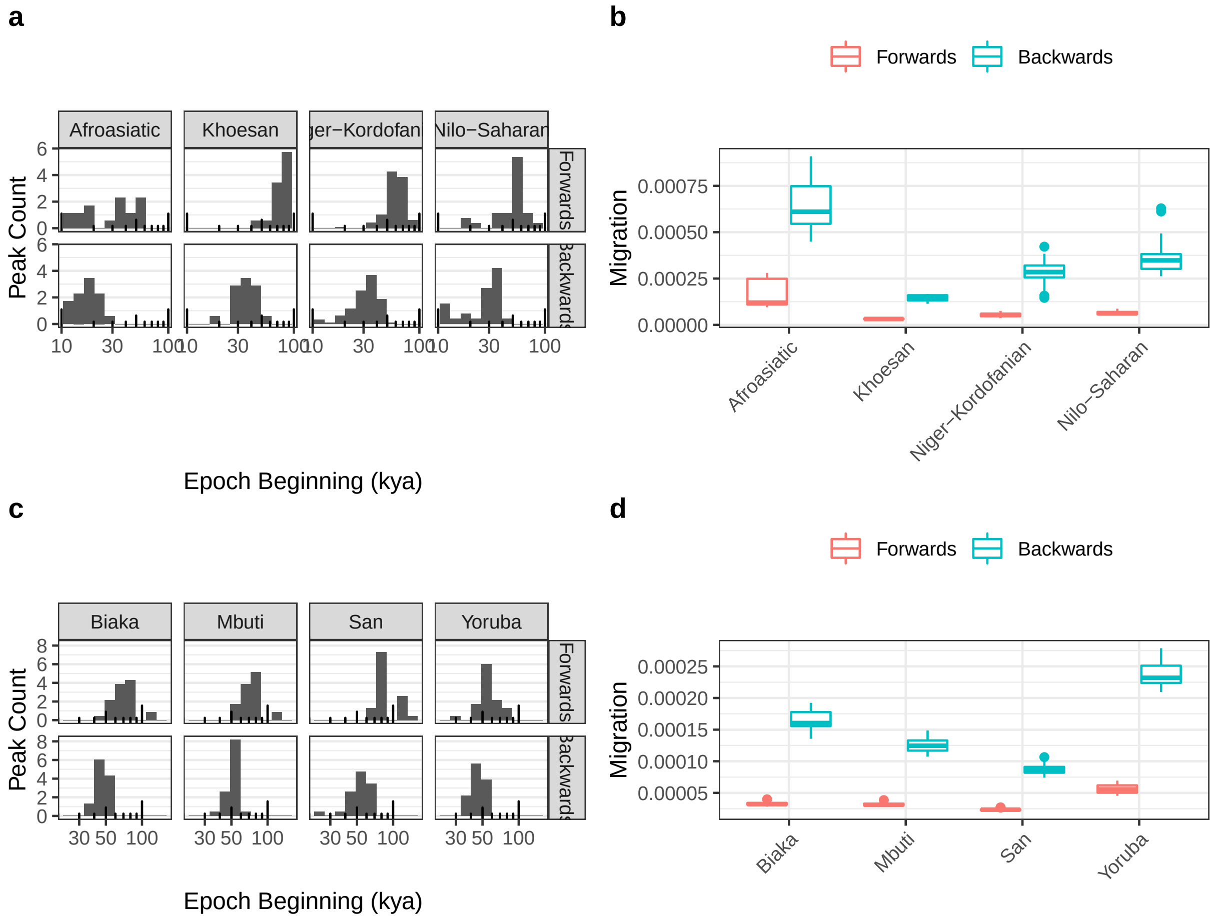

Figure S1: **Timing and average maximum rate of directional migration in HGDP and SGDP. a** Migration is inferred in evenly spaced epochs on the log scale from 3.8 thousand to 3.8 million years ago. For each population in the SGDP, we record the epoch with the highest inferred directional migration rate (the "peak" of migration) and plot this as a histogram. Backwards migration refers to migration from Eurasians to Africans, whilst forward represents the reverse. **b** In the epochs of highest migration identified in a., we record the inferred rate per population and plot these as a boxplot. Whiskers represent 1.5 times the interquartile range. The migration rate is given in proportion of the population replaced per generation. **c** and **d** represent the same analyses as in a. and b. calculated for the Human Genome Diversity Panel, rather than the SGDP.

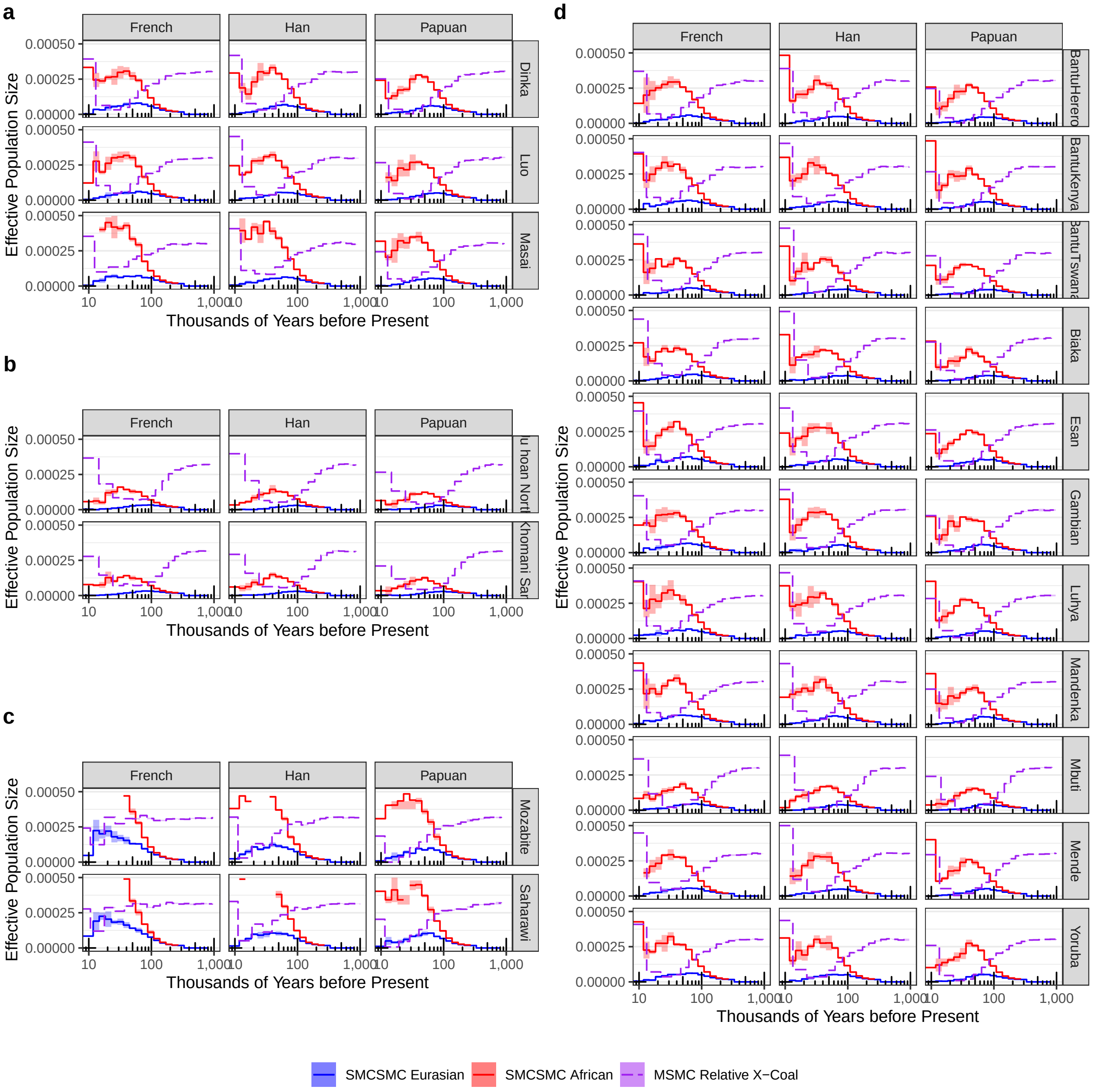

Figure S2: **Inference of directional migration in the Simons Genome Diversity Project.** The `SMCSMC` particle filter was used to infer directional migration rates in both directions from one of three Eurasian populations (French, Han, and Papuan) to one of 18 African populations. 5000 particles were used to approximate the ancestral recombination graph with 10 iterations of variational Bayesian inference to update demographic parameter values. Panels represent a. Nilo-Saharan, b. KhoeSan, c. Afroasiatic, and d. Niger-Kordofanian language families. Alongside the `SMCSMC` inference, we use `MSMC2` to infer the relative cross coalescent rate (RCCR) with default settings and 20 iterations for convergence.

**a**

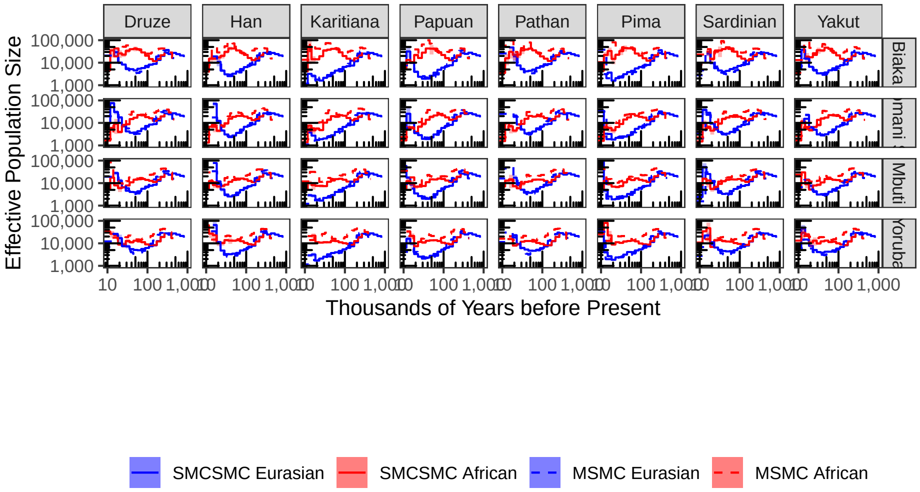

**b**

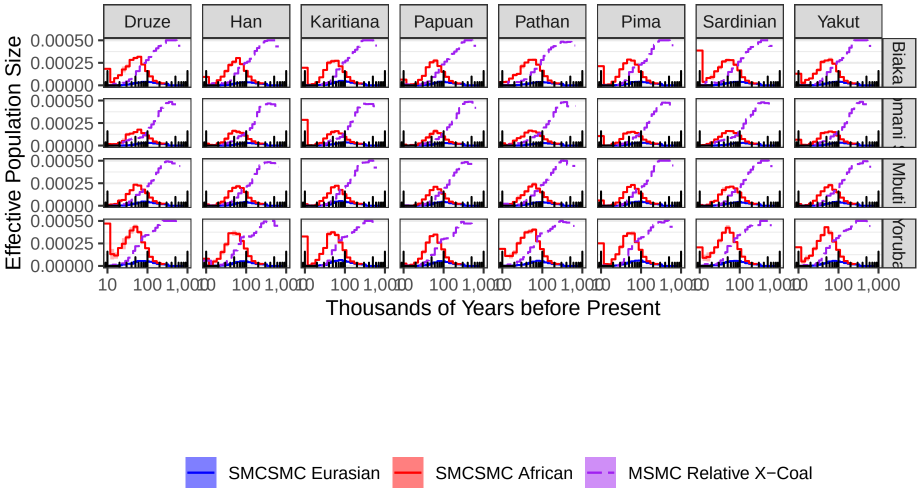

Figure S3: **Demographic inference in a matched subset of the Simons Genome Diversity Panel.** **a.** `SMCSMC` and `MSMC2` inferred effective population size of several populations in the Simons Genome Diversity Panel. These samples were selected to match, as closely as possible, those in the physically phased subet of the Human Genome Diversity Project panel. b. Inferred migration using `SMCSMC` in the Simons Genome Diversity Panel along with the scaled relative cross-coalescent rate estimated by `MSMC2`. 10,000 particles were used to approximate the ancestral recombination graph in the `SMCSMC` particle filter and 25 iterations were used to update demographic parmaeters. 20 iterations were used for `MSMC2`.

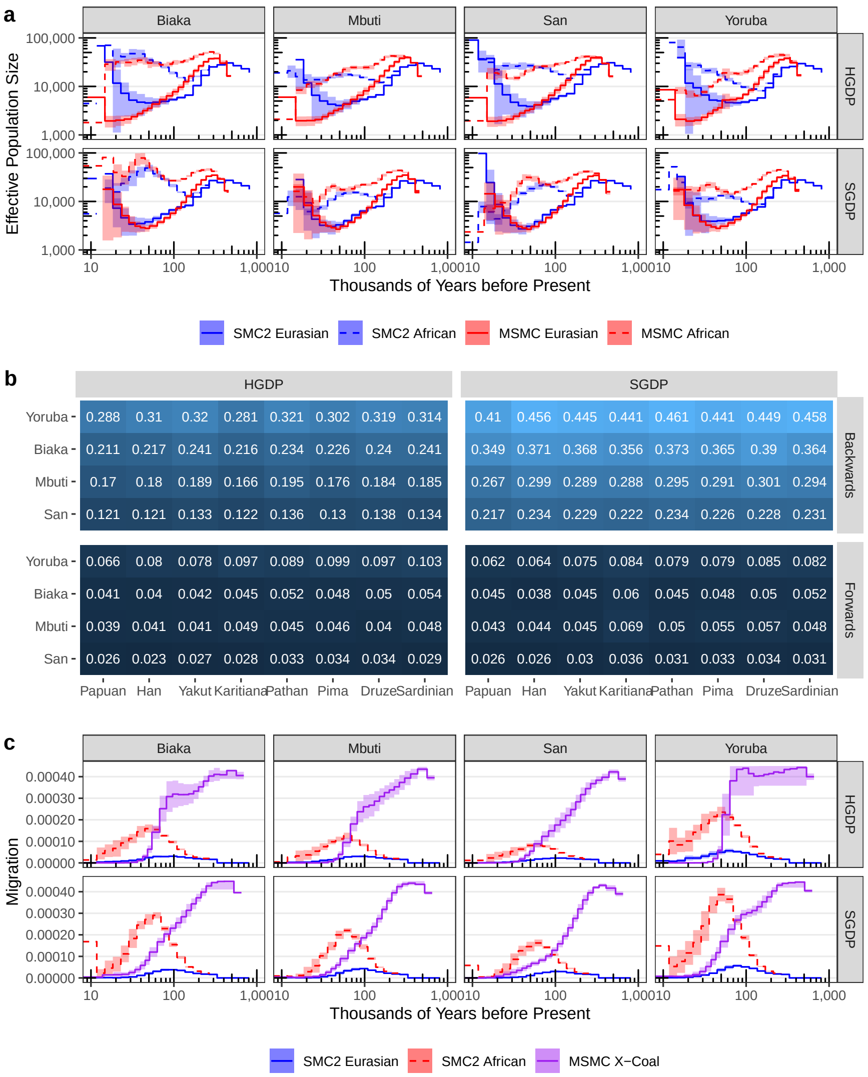

Figure S4: **Inference of directional migration is comparable between data sets and phasing strategies.** We used SMCSMC as in S3 to simultaneously infer directional migration rates and effective population size in the 36 genome physically phased subset of the Human Genome Diversity Panel. We match these 36 genomes with comparable individuals in the Simons Genome Diversity Panel (with the exception of one Papuan population, which has no comparable population in the SGDP) and perform an identical analysis. **a.** Average $N_e$ estimate across four populations in the physically phased subset of the HGDP and the subset of SGDP used to compare with HGDP inference. Inference of population size is averaged over eight Eurasian populations, with the bars representing standard deviation. For MSMC2, the time indexes were averaged to have consistent start and stop times for the steps. **b.** Inferred integrated migration fraction (IMF) from Africa to Eurasians (forwards) and from Eursaians to Africans (backwards) between 40 and 70 kya (see Methods). **c.** Directional migration inference in African populations averaged over Eurasian partners in the two data sets. Shaded regions denote standard deviations. For MSMC, the time indexes were averaged to have consistent start and stop times for plotting.

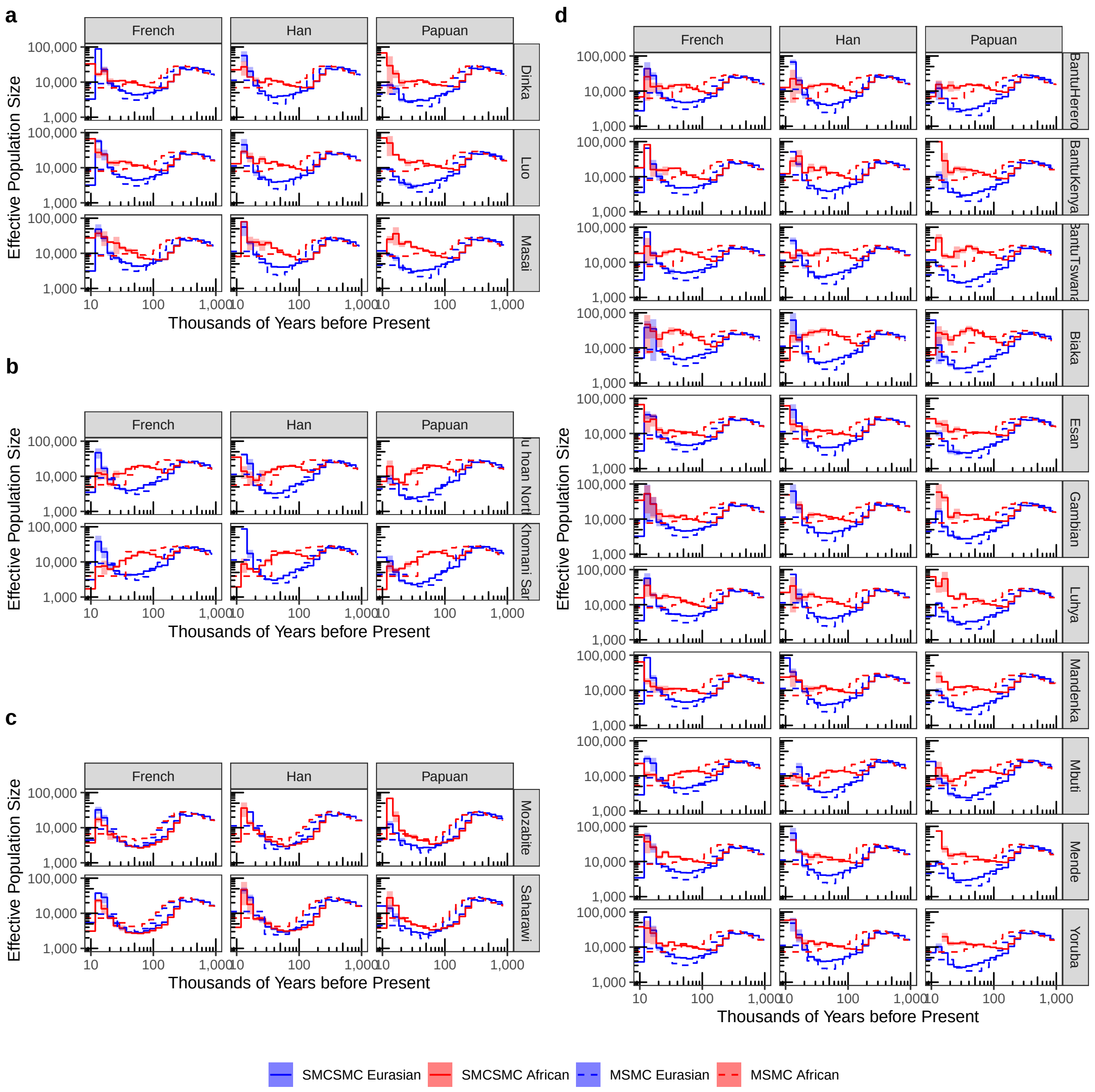

Figure S5: **Inference of historical effective population size in the Simons Genome Diversity Project.** The SMCSMC particle filter was used to infer directional migration rates and effective population size in both directions from one of three Eurasian populations (French, Han, and Papuan) to one of 18 African populations. 5000 particles were used to approximate the ancestral recombination graph with 10 iterations of variational Bayesian inference to update demographic parameter values. Panels represent a. Nilo-Saharan, b. KhoeSan, c. Afroasiatic, and d. Niger-Kordofanian language families. Alongside the SMCSMC inference, we use MSMC2 to infer the same values with default settings and 20 iterations for convergence.

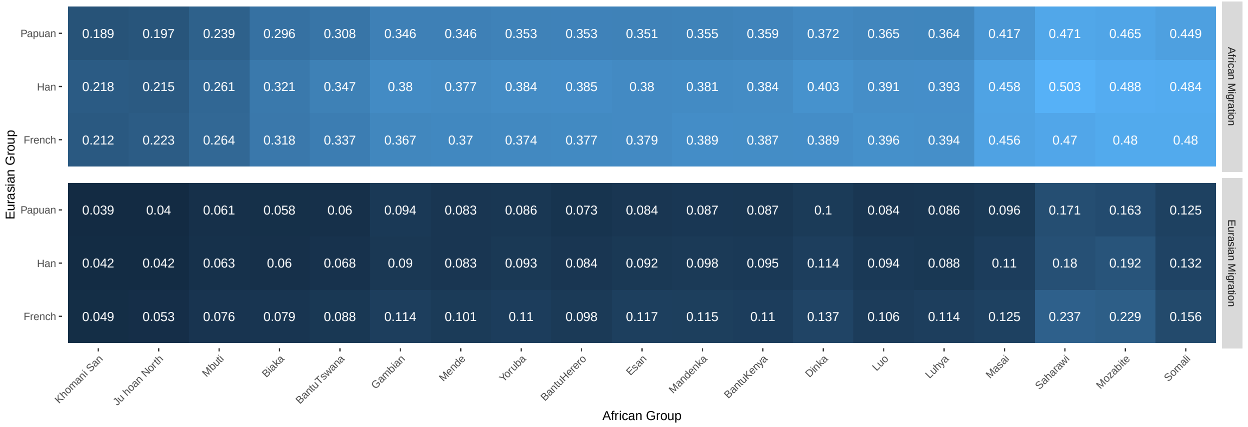

Figure S6: **Integrated migration fraction 40–70kya in SMCSMC analysed SGDP populations.** Following the analysis in Supplemental Section S3, directional migration was integrated by finding the cumulative probability of an individual migrating during the specified epoch. Directional migration backwards from Eurasia to Africa and forwards from Africa to Eurasia (both forward in time) are reported separately. Displayed is the average values from three technical replicates.

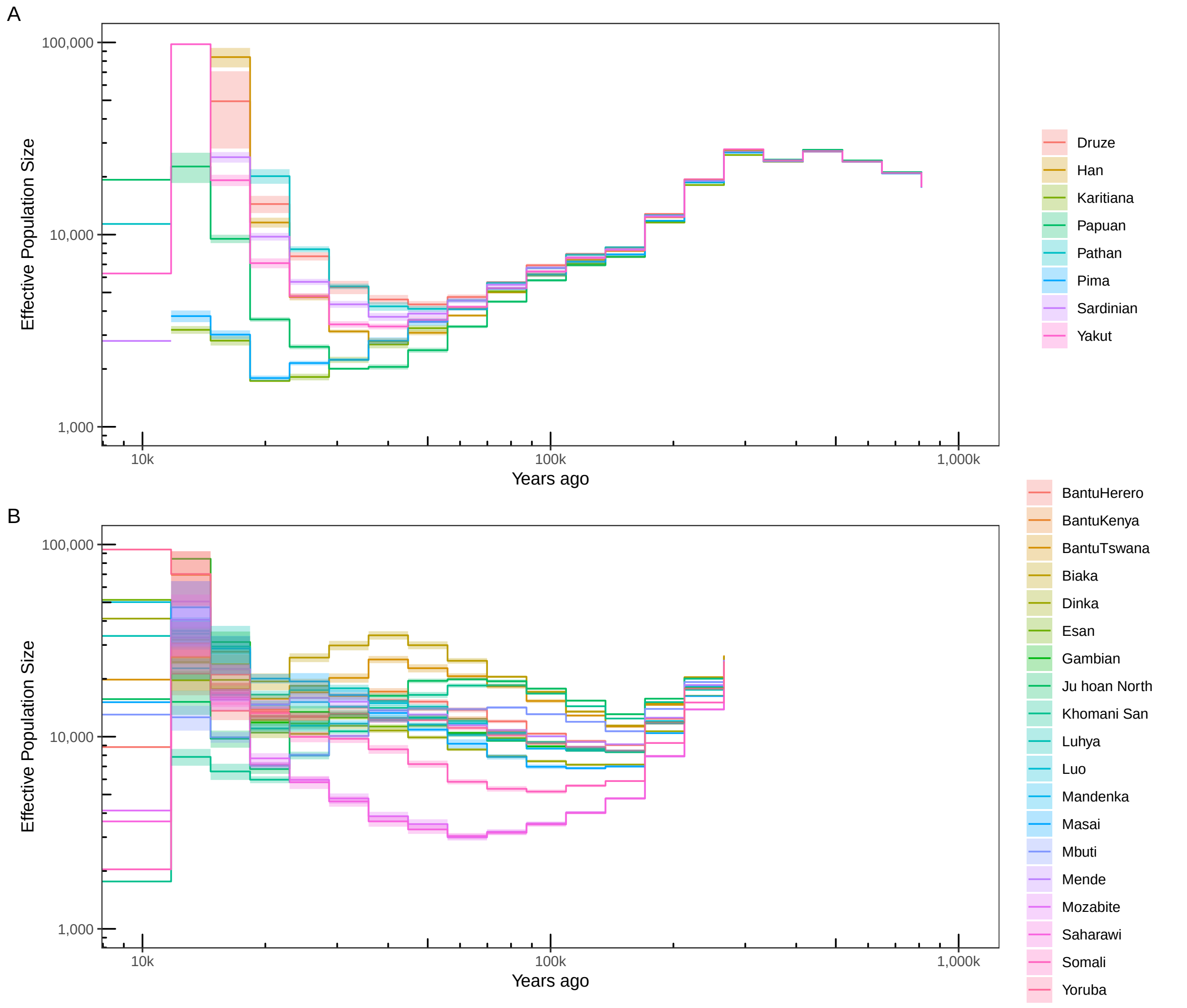

Figure S7: **Estimates of individual population sizes incorporating directional migration.** Using `SMCSMC` the effective population size of global populations in the Simons Genome Diversity Panel is inferred while simultaneously fitting directional migration estimates. Averages are plotted by epoch, with shaded regions denoting the standard deviation. **a.** Estimates of Eurasian population sizes when averaged over Eurasian donor populations. This analysis uses the eight Eurasian populations matched to HGDP populations averaged over the four matched African populations. B. Estimates of African population sizes when averaged over Eurasian recipient populations. This analysis uses the three donor Eurasian populations used for the majority of the analyses in the main text (French, Han, and Papuan) along with the given African populations. Before approximately 250kya, the populations share the same population size within the model, and are not plotted. 10,000 particles are used to approximate the ancestral recombination graph in the SMCSMC particle filter and 15 iterations are used to update demographic parameter values.

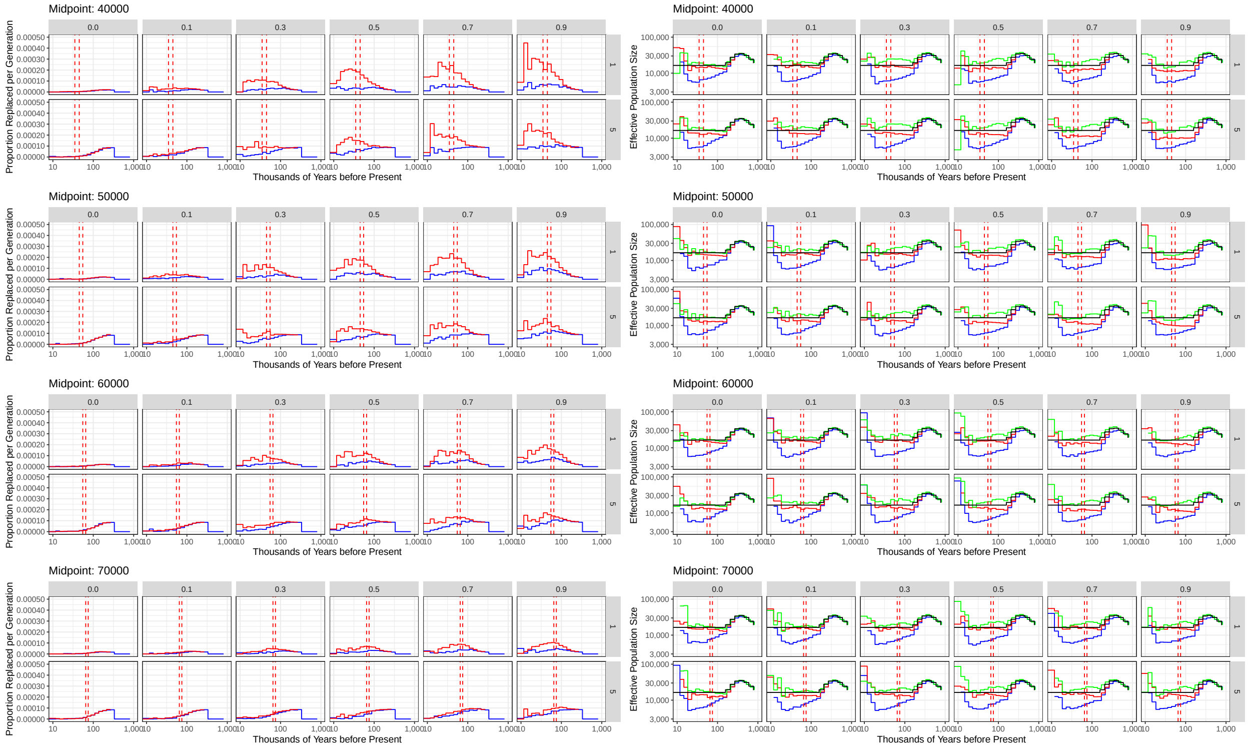

Figure S8: **Simulated migration backwards from effectively Eurasian to effectively African populations.** One gigabase of simulated sequence data was generated with SCRM for two diploid individuals from different populations using a 100kb sliding window approximation of the coalescent with recombination and a demographic model similar to Eurasians and Africans specified in Supplemental section S5. The timing, representing the midpoint of a 10ky migration episode, and the integrated migration fraction (IMF) were systematically varied. In this scenario, only backwards migration was simulated. The left panel displays the recovered directional migration, with red lines representing migration backwards from Eurasia and blue representing migration forwards to Eurasia. The timing of the migration episode is demarcated with dotted red lines. The right panel displays the inferred effective population size of both populations. Additionally, the effective population size of the African-like population was modelled separately, without simultaneously inferring migration. 5000 particles were used in the SMCSMC particle filter to approximate the ancestral recombination graph and 5 iterations of variational Bayesian inference were used to updated demographic parameter values.

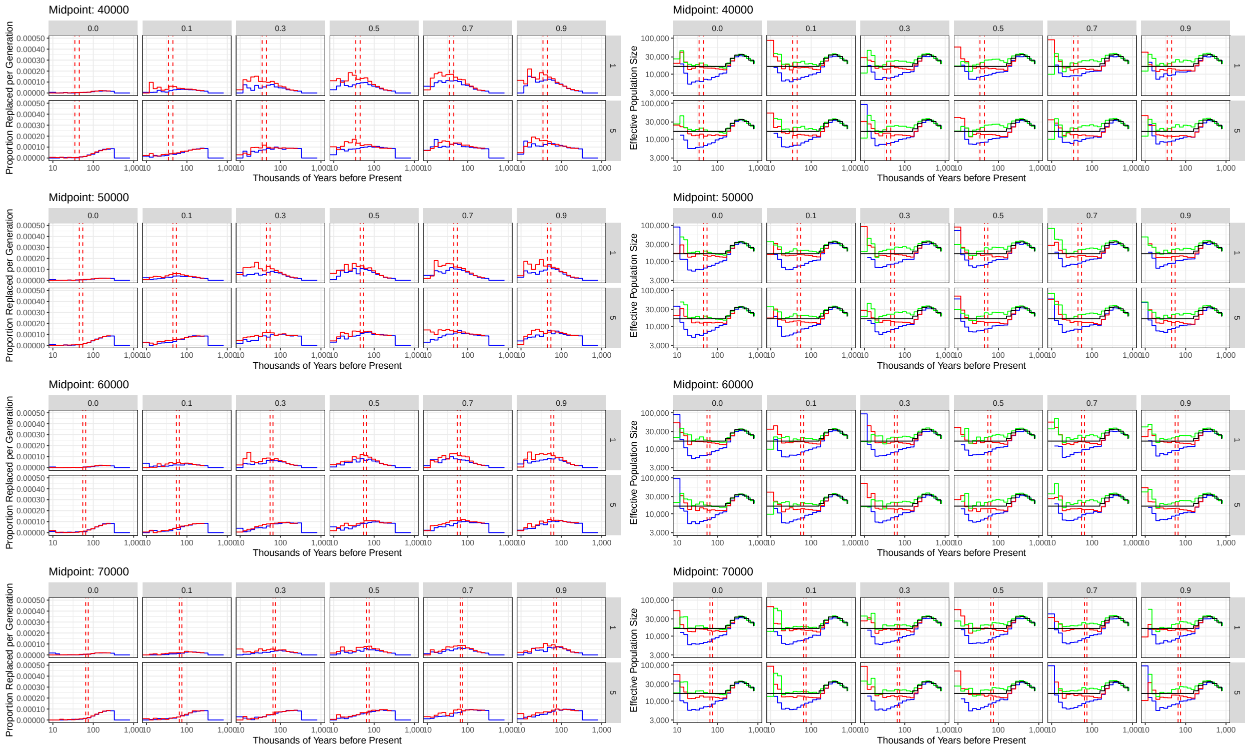

Figure S9: **Simulated bidirectional migration between effectively Eurasian and effectively African populations.** One gigabase of simulated sequence data was generated with `SCRM` for two diploid individuals from different populations using a 100kb sliding window approximation of the coalescent with recombination and a demographic model similar to Eurasians and Africans specified in Supplemental section S5. The timing, representing the midpoint of a 10ky migration episode, and the integrated migration fraction (IMF) were systematically varied. In this scenario, migration between these two populations was simulated with equal rates. The left panel displays the recovered directional migration, with red lines representing migration backwards from Eurasia and blue representing migration forwards to Eurasia. The timing of the migration episode is demarcated with dotted red lines. The right panel displays the inferred effective population size of both populations. Additionally, the effective population size of the African-like population was modelled separately, without simultaneously inferring migration. 5000 particles were used in the `SMCSMC` particle filter to approximate the ancestral recombination graph and 5 iterations of variational Bayesian inference were used to updated demographic parameter values.

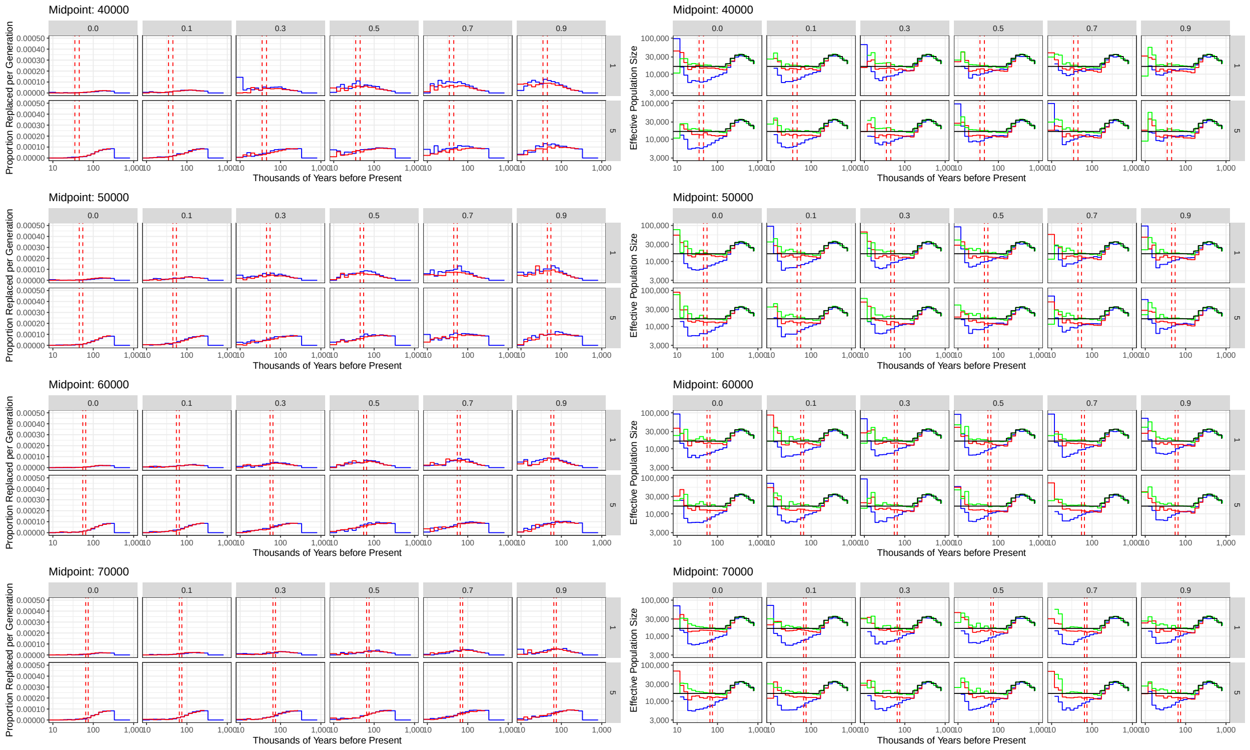

Figure S10: **Simulated migration forwards from effectively African to effectively Eurasian populations.** One gigabase of simulated sequence data was generated with `SCRM` for two diploid individuals from different populations using a 100kb sliding window approximation of the coalescent with recombination and a demographic model similar to Eurasians and Africans specified in Supplemental section S5. The timing, representing the midpoint of a 10ky migration episode, and the integrated migration fraction (IMF) were systematically varied. In this scenario, only forwards migration was simulated. The left panel displays the recovered directional migration, with red lines representing migration backwards from Eurasia and blue representing migration forwards to Eurasia. The timing of the migration episode is demarcated with dotted red lines. The right panel displays the inferred effective population size of both populations. Additionally, the effective population size of the African-like population was modelled separately, without simultaneously inferring migration. 5000 particles were used in the `SMCSMC` particle filter to approximate the ancestral recombination graph and 5 iterations of variational Bayesian inference were used to updated demographic parameter values.

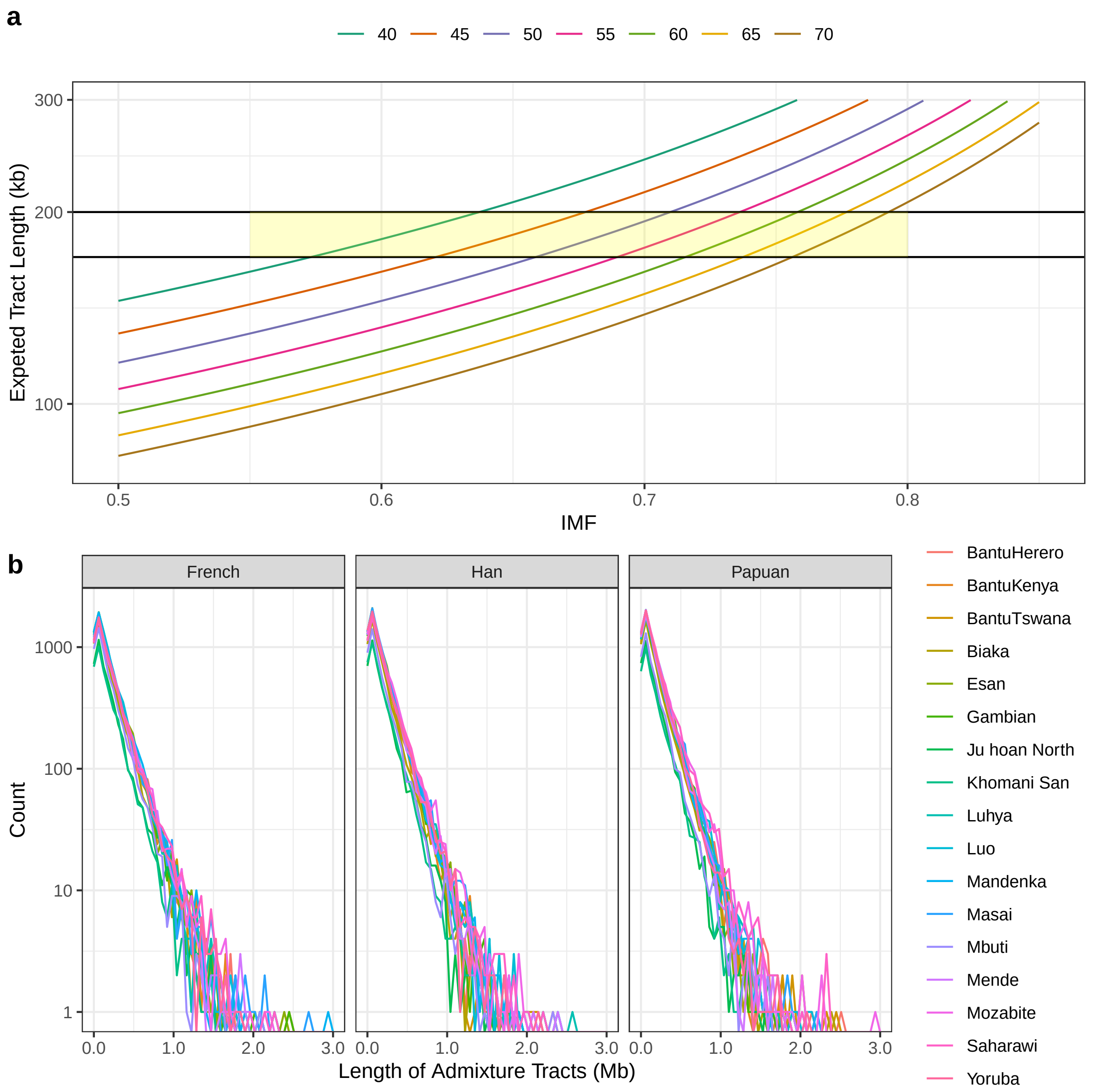

Figure S11: **Analysis of the length of putatively migrated segments**. **a.** Theoretical length distribution of admixture tract rate parameter under varying migration and admixture timing assuming that $L = ((1-m)r(T-1))^{-1}1$, given $L$ length, $m$ symmetrical migration rate, $r$ recombination rate in events per nucleotide, and $T$ time in generations [19]. Shaded region denotes empirical range (with San at the bottom end, and Yoruban at the top end, Supplemental Table S10) of fragments observed in the Simons Genome Diversity Panel. **b.** Following the reconstruction of the ancestral recombination graph using different African and non-African individuals using the SMCSMC particle filter, we use a sample of the posterior distribution of marginal trees to reconstruct putatively migrated segments (see Methods). We plot the length distribution of admixture tracts between individuals in the SGDP using 50 bins. Length is given in megabases (Mb). Isolated from SMCSMC estimated ancestral recombination graphs. Migration rate is given in terms of proportion of the sink population replaced per generation.

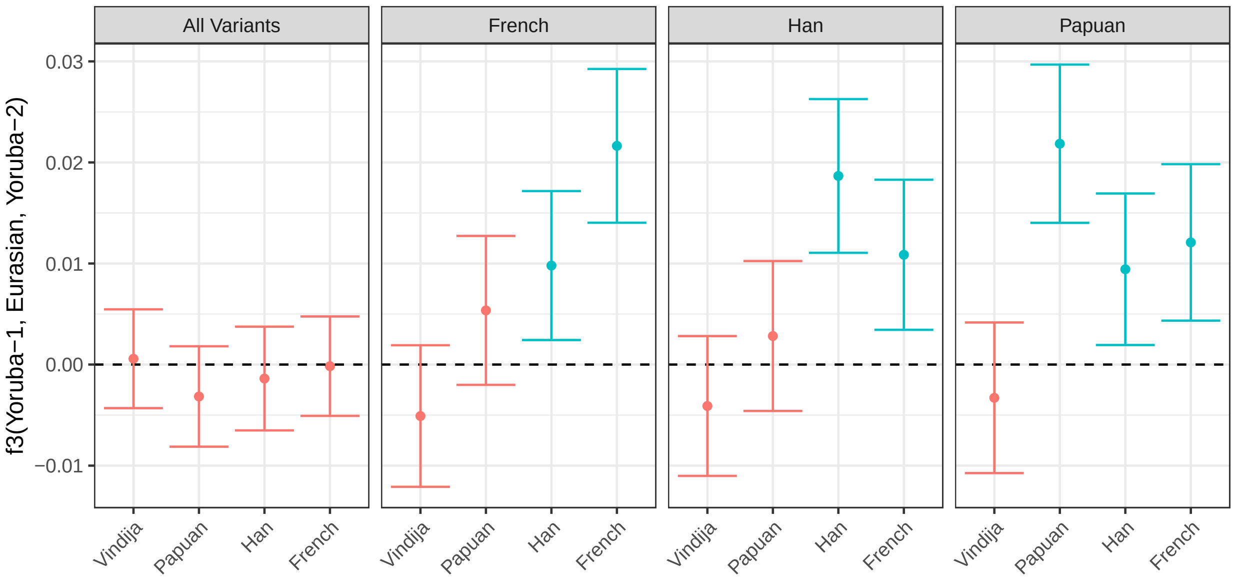

Figure S12: $f_3$ **statistics show evidence for shared drift with Eurasians**. Following the reconstruction of putatively migrated segments from the inferred ancestral recombination graph, we use the Reich Human Origins data set to investigate admixture using drift statistics using `ADMIXTOOLS` and `admixr` (see URLs). Tests are separated into all markers, and those ascertained through `SMCSMC` runs with the French, Han, and Papuans as a comparison Eurasian group. The $f_3$ statistics estimate is plotted, along with 95% C.I. computed via a block jackknife. Statistics which are significantly larger than zero are coloured blue, indicating shared drift.

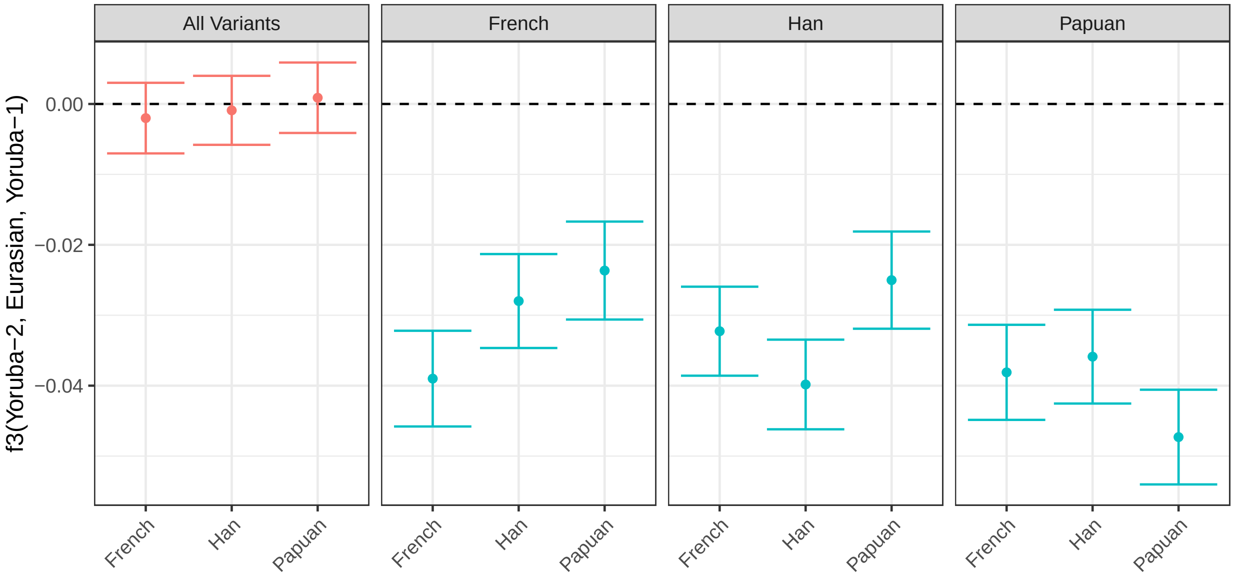

Figure S13: $f_3$ **statistics show evidence for Eurasian admixture**. Following the reconstruction of putatively migrated segments from the inferred ancestral recombination graph, we use the Reich Human Origins data set to investigate admixture using drift statistics using `ADMIXTOOLS` and `admixr` (see URLs). Tests are separated into all markers, and those ascertained through `SMCSMC` runs with the French, Han, and Papuans as a comparison Eurasian group. The $f_3$ statistics estimate is plotted, along with 95% C.I. computed via a block jackknife. Statistics which are significantly less than zero indicate statistical evidence for admixture, and are coloured blue.

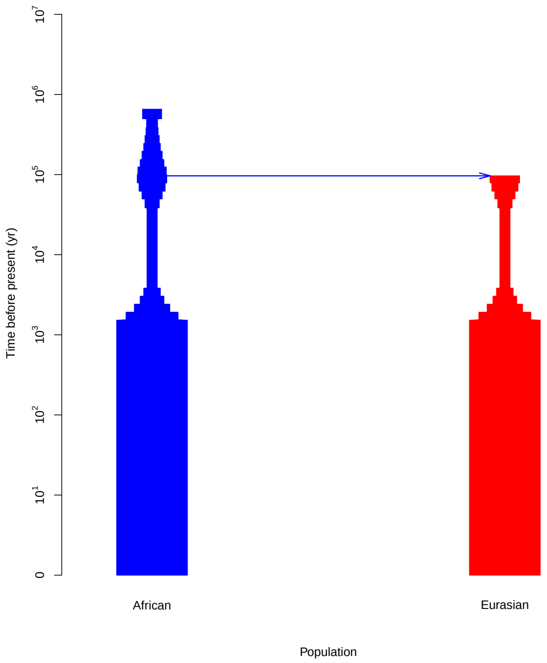

Figure S14: Demographic model used as initialisation for `SMCSMC` analysis visualised using PopDemog [46].

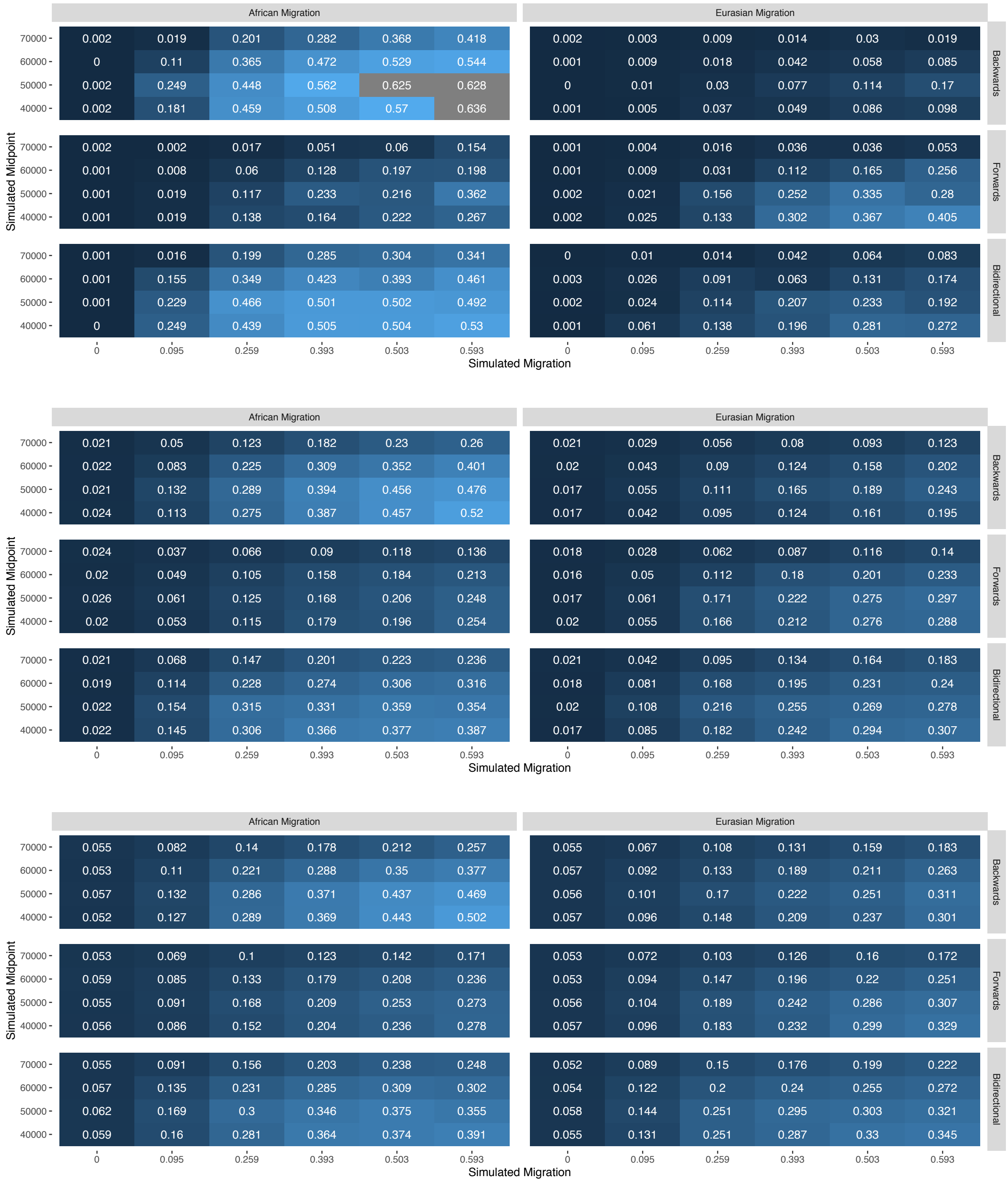

Figure S15: **Integrated migration fraction (IMF) in the last 100ky for three cases of simulated demography shown in Figures S8, S9, and S10.** Simulations were performed as per Supplemental Section S5 with an additional parameter for the initial migration rate used to initialise the SMCSMC particle filter. The timing and IMF simulated are as per the aforementioned figures. From top to bottom, inference was initiated with 0, 1, and 5 $4N_0$ population replacement per generation in the specified direction (backwards, bidirectionally, and forwards).

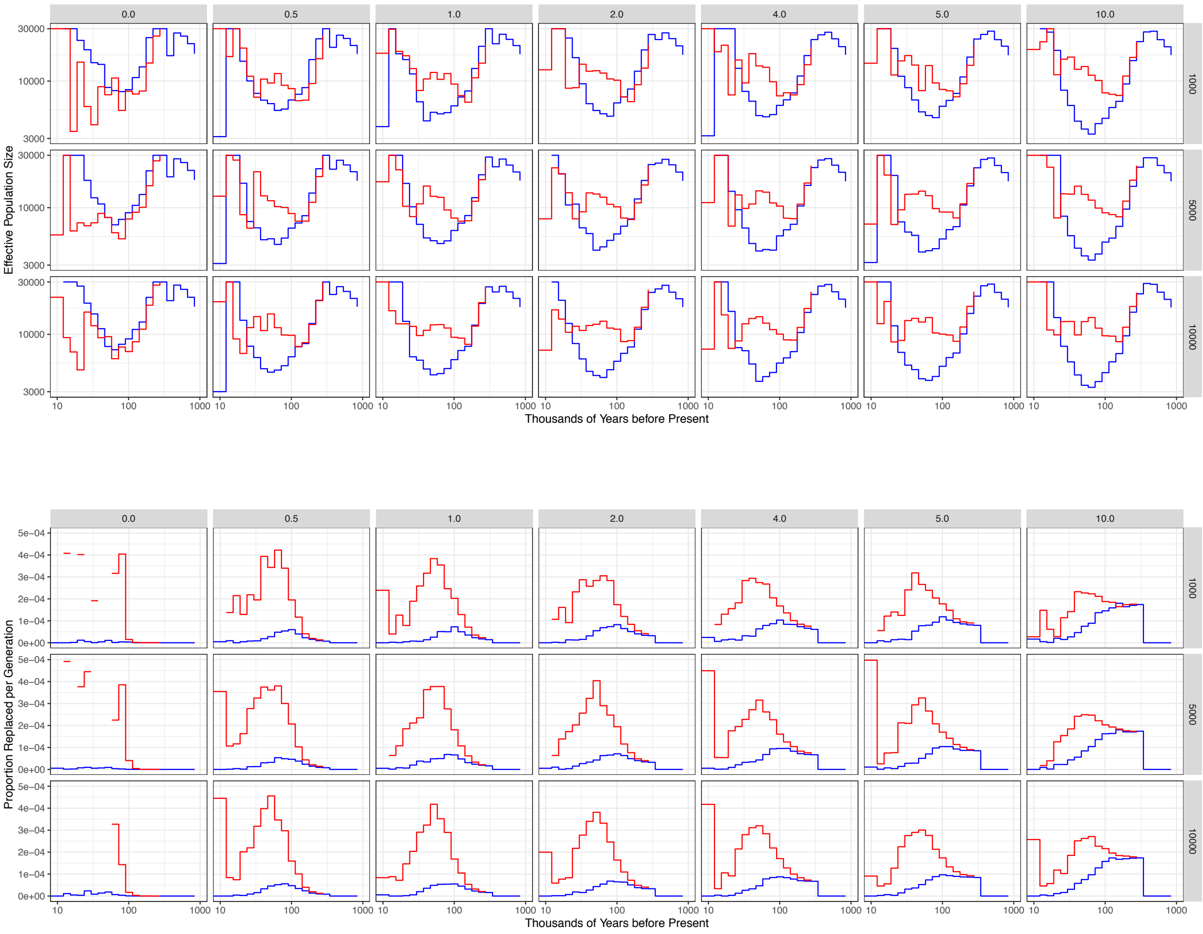

Figure S16: **The effect of initial migration parameter on demographic inference.** Effective population size and migration history of a Yoruban (S_Yoruba-1) and a French (S_French-1) individual from the Simons Genome Diversity panel were modelled with SMCSMC. The initial migration proportion, in units of 4 $N_0$ proportion of the population replaced per generation was varied along the X axis, while the number of particles is varied along the Y. 10 iterations of variational Bayes was used for parameter inference, while 5000 particles were used infer the ancestral recombination graph.

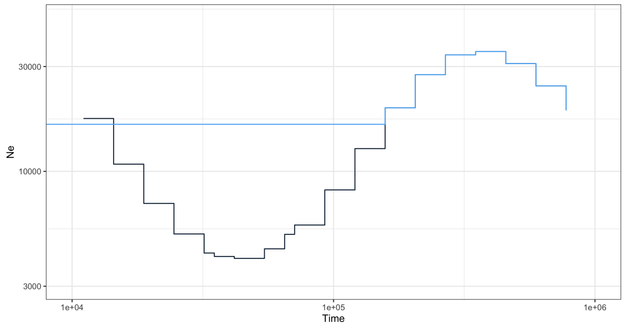

Figure S17: **Effective Population size model used for simulations.** Following the simulation procedure in Supplemental Section S5, the population size model is plotted per epoch, with the effectively African population plotted in blue and the effectively non-African population in black.

#### S2 Supplemental Tables

| Language Family | Partner Population | Mean AFR IMF (SD) | Mean EUR IMF (SD) |
|---|---|---|---|
| Afroasiatic | All | 0.477(0.015) | 0.176(0.038) |
| Afroasiatic | French | 0.477(0.006) | 0.207(0.044) |
| Afroasiatic | Han | 0.492(0.01) | 0.168(0.032) |
| Afroasiatic | Papuan | 0.462(0.011) | 0.153(0.024) |
| Khoesan | All | 0.209(0.013) | 0.044(0.006) |
| Khoesan | French | 0.217(0.007) | 0.051(0.003) |
| Khoesan | Han | 0.217(0.002) | 0.042(0) |
| Khoesan | Papuan | 0.193(0.006) | 0.04(0) |
| Niger-Kordofanian | All | 0.352(0.04) | 0.088(0.017) |
| Niger-Kordofanian | French | 0.36(0.039) | 0.102(0.015) |
| Niger-Kordofanian | Han | 0.363(0.04) | 0.083(0.013) |
| Niger-Kordofanian | Papuan | 0.334(0.038) | 0.078(0.013) |
| Nilo-Saharan | All | 0.405(0.033) | 0.107(0.016) |
| Nilo-Saharan | French | 0.414(0.037) | 0.123(0.016) |
| Nilo-Saharan | Han | 0.417(0.036) | 0.106(0.01) |
| Nilo-Saharan | Papuan | 0.385(0.029) | 0.093(0.008) |

Table S1: **Integrated Migration Fraction (IMF) in either direction averaged over African language groups.** `SMCSMC` was used as in Supplemental Section S3 to infer directional migration and effective population size between populations in the Simons Genome Diversity Project. The total migration between each African and non-African population was integrated in the epoch 30–70kya (see Methods) and averaged over language family. Abbreviations: Integrated Migration Fraction (IMF), Stanard Deviation (SD), EUR (Eurasian), AFR (African)

| | Estimate | Std. Error | t value | Pr(>\|t\|) |
|---|---|---|---|---|
| (Intercept) | 0.3807 | 0.0036 | 105.42 | 1.79e-47 |
| Papuan | -0.0271 | 0.0017 | -15.60 | 7.44e-18 |
| BantuKenya | 0.0051 | 0.0050 | 1.02 | 3.16e-01 |
| BantuTswana | -0.0410 | 0.0050 | -8.13 | 9.49e-10 |
| Biaka | -0.0598 | 0.0050 | -11.87 | 3.53e-14 |
| Dinka | 0.0160 | 0.0050 | 3.17 | 3.05e-03 |
| Esan | -0.0016 | 0.0050 | -0.32 | 7.51e-01 |
| Gambian | -0.0073 | 0.0050 | -1.44 | 1.58e-01 |
| Ju hoan North | -0.1603 | 0.0050 | -31.79 | 1.82e-28 |
| Khomani San | -0.1653 | 0.0050 | -32.80 | 5.96e-29 |
| Luhya | 0.0118 | 0.0050 | 2.34 | 2.46e-02 |
| Luo | 0.0123 | 0.0050 | 2.45 | 1.94e-02 |
| Mandenka | 0.0032 | 0.0050 | 0.63 | 5.34e-01 |
| Masai | 0.0719 | 0.0050 | 14.25 | 1.32e-16 |
| Mbuti | -0.1171 | 0.0050 | -23.22 | 1.17e-23 |
| Mende | -0.0071 | 0.0050 | -1.42 | 1.65e-01 |
| Mozabite | 0.1060 | 0.0050 | 21.02 | 3.63e-22 |
| Saharawi | 0.1097 | 0.0050 | 21.75 | 1.12e-22 |
| Somali | 0.0995 | 0.0050 | 19.73 | 3.12e-21 |
| Yoruba | -0.0013 | 0.0050 | -0.26 | 7.97e-01 |

Table S2: **Linear model predicting integrated migration fraction in the SGDP.** The integrated migration fraction (IMF) in the epoch 30–70 kya is obtained as per the Methods section in the Simons Genome Diversity Project. A binary variable representing Papuan / not Papuan Eurasian donor and categorical variable representing African population were used to predict the IMF in a simple linear model. When adjusted for the different African populations, Papuans contribute less IMF than do other Eurasian partners (French and Han).

| | Estimate | Std. Error | t value | Pr(>\|t\|) |
|---|---|---|---|---|
| (Intercept) | 0.3392 | 0.0037 | 92.92 | 1.72e-39 |
| Papuan | -0.0252 | 0.0042 | -5.95 | 1.43e-06 |
| San | -0.1942 | 0.0050 | -38.94 | 6.85e-28 |
| Mbuti | -0.1410 | 0.0050 | -28.27 | 1.07e-23 |
| Biaka | -0.0883 | 0.0050 | -17.69 | 9.33e-18 |

Table S3: **Linear model predicting integrated migration fraction in the HGDP.** The integrated migration fraction (IMF) in the epoch 30–70 kya is obtained as per the Methods section in the physically phased subset of the Human Genome Diversity Project. A binary variable representing Papuan / not Papuan Eurasian donor and categorical variable representing African population were used to predict the IMF in a simple linear model. When adjusted for the different African populations, Papuans contribute less IMF than do other Eurasian partners (French and Han).

| Language Family | Comparison Family | Mean Difference (95% CI) | Adjusted P |
|---|---|---|---|
| Niger-Kordofanian | Nilo-Saharan | -0.053 (-0.081–0.025) | 1.11e-03 |
| Niger-Kordofanian | Khoesan | 0.143 (0.125-0.161) | 4.39e-14 |
| Niger-Kordofanian | Afroasiatic | -0.125 (-0.142–0.107) | 1.67e-15 |
| Nilo-Saharan | Niger-Kordofanian | 0.053 (0.025-0.081) | 1.11e-03 |
| Nilo-Saharan | Khoesan | 0.196 (0.169-0.223) | 6.91e-09 |
| Nilo-Saharan | Afroasiatic | -0.072 (-0.098–0.045) | 1.23e-04 |
| Khoesan | Niger-Kordofanian | -0.143 (-0.161–0.125) | 4.39e-14 |
| Khoesan | Nilo-Saharan | -0.196 (-0.223–0.169) | 6.91e-09 |
| Khoesan | Afroasiatic | -0.268 (-0.284–0.252) | 3.62e-13 |
| Afroasiatic | Niger-Kordofanian | 0.125 (0.107-0.142) | 1.67e-15 |
| Afroasiatic | Nilo-Saharan | 0.072 (0.045-0.098) | 1.23e-04 |
| Afroasiatic | Khoesan | 0.268 (0.252-0.284) | 3.62e-13 |
| CAHG | Niger-Kordofanian | -0.084 (-0.119–0.049) | 1.18e-03 |

Table S4: **Differences in African IMF in the SGDP.** The integrated migration fraction (IMF) in the epoch 30–70kya is calculated as per the Methods section for all comparisons in the Simons Genome Diversity Project (SGDP), and a two tailed *t*-test is used to statistically test for differences between the inferred migration in African language groups. Averaged over three technical replicates to account for the influence of stochastic sampling variation. P values corrected for multiple testing using the Bonferroni method. Abbreviations: CAHG, Central African Hunter-Gatherers (include Mbuti and Biaka); CI, Confidence Interval.

| Statistic | D | Z |
|---|---|---|
| D(KhomaniSan-1, Yoruba-1, Yoruba-2, Chimp) | -0.181 | -28.403 |
| D(Mbuti-1, Yoruba-1, Yoruba-2, Chimp) | -0.135 | -19.554 |
| D(Papuan-1, Yoruba-1, Yoruba-2, Chimp) | -0.026 | -3.422 |
| D(French-1, Yoruba-1, Yoruba-2, Chimp) | -0.006 | -0.866 |
| D(Han-1, Yoruba-1, Yoruba-2, Chimp) | 0.001 | 0.072 |
| D(KhomaniSan-1, Yoruba-2, Yoruba-1, Chimp) | -0.187 | -28.109 |
| D(Mbuti-1, Yoruba-2, Yoruba-1, Chimp) | -0.130 | -19.323 |
| D(Papuan-1, Yoruba-2, Yoruba-1, Chimp) | -0.008 | -1.003 |
| D(French-1, Yoruba-2, Yoruba-1, Chimp) | 0.030 | 4.355 |
| D(Han-1, Yoruba-2, Yoruba-1, Chimp) | 0.056 | 8.037 |

Table S5: Putatively migrated segments of a Yoruban are closer to Out of Africa groups than a comparable Yoruban.

| Statistic | D | Z |
|---|---|---|
| D(Mbuti-1, Yoruba-1, Vindija, Chimp) | -0.001 | -0.141 |
| D(Mbuti-1, Yoruba-2, Vindija, Chimp) | -0.003 | -0.306 |

Table S6: No difference in allele sharing with Vindija Neanderthal over Mbuti baseline.

| Statistic | D | Z |
|---|---|---|
| D(Yoruba-2, Yoruba-1, Vindija, Chimp) | 0.000 | 0.012 |

Table S7: No difference in allele sharing with Vindija Neanderthal.

| Statistic | D | Z |
|---|---|---|
| D(Vindija, Altai, Yoruba-1, Chimp) | 0.024 | 1.095 |
| D(Vindija, Altai, Yoruba-2, Chimp) | 0.034 | 1.526 |
| D(Vindija, Altai, Mbuti-1, Chimp) | 0.002 | 0.103 |
| D(Vindija, Altai, KhomaniSan-1, Chimp) | 0.023 | 1.008 |

Table S8: No increased affinity to Vindija Neanderthal over Altai, as would be expected if the source of any Neanderthal ancestry was Eurasian.

| Name | ID | Source |
|---|---|---|
| French | S_French-1 | SGDP |
| Han | S_Han-1 | SGDP |
| Papuan | S_Papuan-1 | SGDP |
| BantuHerero | S_BantuHerero-1 | SGDP |
| BantuKenya | S_BantuKenya-1 | SGDP |
| BantuTswana | S_BantuTswana-1 | SGDP |
| Biaka | S_Biaka-1 | SGDP |
| Dinka | B_Dinka-3 | SGDP |
| Esan | S_Esan-1 | SGDP |
| Gambian | S_Gambian-1 | SGDP |
| Ju hoan North | S_Ju_hoan_North-1 | SGDP |
| Khomani San | S_Khomani_San-1 | SGDP |
| Luhya | S_Luhya-1 | SGDP |
| Luo | S_Luo-1 | SGDP |
| Mandenka | S_Mandenka-1 | SGDP |
| Masai | S_Masai-1 | SGDP |
| Mbuti | S_Mbuti-1 | SGDP |
| Mende | S_Mende-1 | SGDP |
| Mozabite | S_Mozabite-1 | SGDP |
| Saharawi | S_Saharawi-1 | SGDP |
| Somali | S_Somali-1 | SGDP |
| Yoruba | S_Yoruba-1 | SGDP |
| Druze | HGDP00562 | HGDP |
| Han | HGDP00774 | HGDP |
| Karitiana | HGDP01013 | HGDP |
| PapuanHighlands | HGDP00549 | HGDP |
| PapuanSepik | HGDP00542 | HGDP |
| Pathan | HGDP00224 | HGDP |
| Pima | HGDP01043 | HGDP |
| Sardinian | HGDP00670 | HGDP |
| Yakut | HGDP00946 | HGDP |
| Yoruba | HGDP00930 | HGDP |
| San | HGDP01029 | HGDP |
| Mbuti | HGDP00450 | HGDP |
| Biaka | HGDP00460 | HGDP |
| Vindija | Vindija.DG | Pruefer et al. 2017 |
| Altai | Altai_published.DG | Pruefer et al. 2013 |
| Denisovan | Deniosva_published.DG | Myers et al 2012 |

Table S9: Sample IDs of the individuals used in this article and relevant resources. SGDP = Simons Genome Diversity Project, HGDP = Human Genome Diversity Panel.

| African Population | Mean (SD) | Total (Mb) | Mean (SD) | Total (Mb) | Mean (SD) | Total (Mb) |
|---|---|---|---|---|---|---|
| Mbuti | 172.204 (189.255) | 939.371 | 165.822 (182.549) | 850.335 | 163.352 (181.607) | 756.320 |
| Biaka | 172.332 (184.183) | 1057.083 | 169.296 (185.457) | 1052.004 | 170.149 (185.853) | 1044.036 |
| Khomani San | 173.06 (188.492) | 671.125 | 169.005 (183.871) | 706.777 | 166.33 (175.899) | 597.125 |
| Ju hoan North | 175.415 (195.979) | 747.618 | 171.909 (187.835) | 711.186 | 161.146 (175.476) | 656.024 |
| Luhya | 177.303 (193.164) | 1370.729 | 181.025 (197.94) | 1403.123 | 175.821 (192.259) | 1211.762 |
| Esan | 177.483 (191.148) | 1364.132 | 178.429 (190.819) | 1426.180 | 170.451 (183.886) | 1270.204 |
| Gambian | 177.804 (195.646) | 1333.529 | 177.099 (191.542) | 1304.155 | 173.892 (185.544) | 1213.071 |
| Luo | 178.653 (192.447) | 1368.481 | 175.618 (192.833) | 1361.917 | 180.635 (201.159) | 1348.798 |
| BantuHerero | 179.691 (194.451) | 1354.869 | 176.473 (188.855) | 1327.779 | 178.544 (198.496) | 1310.511 |
| BantuTswana | 180.309 (195.176) | 1150.551 | 174.826 (188.153) | 1159.796 | 172.542 (192.6) | 1096.674 |
| Mende | 182.087 (200.591) | 1256.580 | 177.24 (197.23) | 1288.533 | 169.901 (182.944) | 1191.515 |
| Yoruba | 184.255 (199.949) | 1283.151 | 176.276 (193.69) | 1361.031 | 169.867 (181.948) | 1275.701 |
| BantuKenya | 184.419 (200.395) | 1265.297 | 173.485 (191.175) | 1269.739 | 179.415 (189.784) | 1214.104 |
| Mandenka | 185.35 (206.972) | 1411.440 | 176.908 (194.337) | 1328.754 | 172.03 (182.984) | 1265.625 |
| Masai | 186.769 (209.675) | 1353.890 | 182.366 (198.06) | 1483.361 | 181.22 (199.255) | 1438.160 |
| Mozabite | 194.378 (214.029) | 1318.273 | 193.469 (211.945) | 1557.040 | 188.867 (210.994) | 1532.842 |
| Saharawi | 197.668 (218.662) | 1383.280 | 196.156 (217.233) | 1406.245 | 195.045 (217.263) | 1585.907 |

Table S10: Summary of the length distribution for putatively migrated segments in different African individuals. Means and standard deviations are given in kilobases (kb) while the total length of all segments is given in megabases (Mb).

| Statistic | D | Z |
|---|---|---|
| D(KhomaniSan-1, Yoruba-1, French-1, Chimp) | -0.173 | -25.685 |
| D(KhomaniSan-1, Yoruba-1, Han-1, Chimp) | -0.208 | -29.150 |
| D(KhomaniSan-1, Yoruba-1, Papuan-1, Chimp) | -0.174 | -24.085 |
| D(KhomaniSan-1, Yoruba-2, French-1, Chimp) | -0.143 | -19.523 |
| D(KhomaniSan-1, Yoruba-2, Han-1, Chimp) | -0.161 | -22.327 |
| D(KhomaniSan-1, Yoruba-2, Papuan-1, Chimp) | -0.160 | -21.258 |
| D(Mbuti-1, Yoruba-1, French-1, Chimp) | -0.136 | -19.835 |
| D(Mbuti-1, Yoruba-1, Han-1, Chimp) | -0.167 | -23.875 |
| D(Mbuti-1, Yoruba-1, Papuan-1, Chimp) | -0.125 | -17.088 |
| D(Mbuti-1, Yoruba-2, French-1, Chimp) | -0.103 | -14.514 |
| D(Mbuti-1, Yoruba-2, Han-1, Chimp) | -0.116 | -16.935 |
| D(Mbuti-1, Yoruba-2, Papuan-1, Chimp) | -0.109 | -14.459 |

Table S11: Both Yorubans share more alleles with OoA populations than San or Mbuti. The individual used to ascertain segments shares more alleles than a comparable individual.

| Statistic | D | Z |
|---|---|---|
| D(Yoruba-2, Yoruba-1, French-1, Chimp) | -0.036 | -5.122 |
| D(Yoruba-2, Yoruba-1, Han-1, Chimp) | -0.056 | -7.888 |
| D(Yoruba-2, Yoruba-1, Papuan-1, Chimp) | -0.018 | -2.483 |

Table S12: Putatively migrated segments were isolated as per Methods in a Yoruban individual modelled with a French individual. The Yoruban used to ascertain segment is more closely related to OoA groups than a comparable Yoruban, in markers falling in the segments.

| | | Segments | | | All markers | | |
|---|---|---|---|---|---|---|---|
| African | Eurasian | D | Std. Err. | Z | D | Std. Err | Z |
| Khomani San | S_Han-1.DG | 0.130 | 0.010 | 12.666 | -0.002 | 0.006 | -0.423 |
| Ju hoan North | S_Han-1.DG | 0.111 | 0.010 | 11.229 | 0.003 | 0.005 | 0.584 |
| Khomani San | S_Mongola-2.DG | 0.102 | 0.010 | 9.992 | -0.001 | 0.005 | -0.104 |
| Khomani San | S_Adygei-2.DG | 0.100 | 0.010 | 9.605 | 0.003 | 0.006 | 0.568 |
| Khomani San | S_Igorot-1.DG | 0.099 | 0.010 | 9.789 | -0.003 | 0.005 | -0.607 |
| Khomani San | S_Kyrgyz-1.DG | 0.099 | 0.010 | 9.963 | 0.000 | 0.005 | 0.023 |
| Khomani San | S_Uygur-1.DG | 0.098 | 0.010 | 10.161 | 0.002 | 0.005 | 0.287 |
| Khomani San | Kharia.DG | 0.098 | 0.010 | 9.940 | 0.003 | 0.005 | 0.594 |
| Khomani San | Kyrgyz.DG | 0.097 | 0.010 | 10.148 | 0.003 | 0.005 | 0.517 |
| Khomani San | S_Sardinian-1.DG | 0.097 | 0.010 | 9.428 | 0.006 | 0.006 | 1.081 |
| Khomani San | Russia_Scythian_IA_questionable | 0.097 | 0.010 | 9.310 | 0.004 | 0.005 | 0.798 |
| Khomani San | S_Dusun-2.DG | 0.097 | 0.011 | 9.147 | -0.001 | 0.006 | -0.211 |
| Khomani San | S_Tuscan-2.DG | 0.096 | 0.010 | 9.452 | 0.002 | 0.006 | 0.316 |
| Khomani San | S_Even-2.DG | 0.096 | 0.010 | 9.613 | -0.001 | 0.005 | -0.199 |
| Khomani San | S_Igorot-2.DG | 0.096 | 0.010 | 9.401 | 0.002 | 0.005 | 0.366 |
| Khomani San | S_Even-1.DG | 0.096 | 0.010 | 9.355 | 0.000 | 0.005 | 0.043 |
| Khomani San | S_Dai-1.DG | 0.096 | 0.011 | 8.970 | -0.001 | 0.006 | -0.146 |
| Khomani San | Czech_N | 0.096 | 0.010 | 9.674 | 0.001 | 0.006 | 0.204 |
| Khomani San | Ireland_MN.SG | 0.095 | 0.011 | 9.053 | 0.006 | 0.006 | 1.042 |
| Khomani San | Iberia_HG | 0.095 | 0.011 | 8.693 | 0.007 | 0.006 | 1.171 |

Table S13: Top 20 D Statistics for D(Test African, Partner African; Eurasian, Chimp) calculated for variants in putatively migrated segments with more than a combined 10,000 variants. Segments were isolated in a test individual (Sample-1) and compared to a partner individual (Sample-2) from the Simons Genome Diversity Panel. Segments were isolated using a Han Chinese individual to estimate migration, and the populations isolated are representative of Han affinity.

| Statistic | D | Z | Interpretation |
|---|---|---|---|
| D(French-1, Yoruba-2, Yoruba-1, Chimp) | 0.030 | 4.355 | In the subset of variants in putatively migrated segments, there is excessive allele sharing between the test Yoruban and Out of Africa groups (French and Han). |
| D(Han-1, Yoruba-2, Yoruba-1, Chimp) | 0.056 | 8.037 | |
| D(Yoruba-2, Yoruba-1, Vindija, Chimp) | 0.000 | 0.012 | The test Yoruban is no closer to the Vindija Neanderthal than a comparable Yoruban. |
| D(Mbuti-1, Yoruba-1, Vindija, Chimp) | -0.001 | -0.141 | |
| D(Mbuti-1, Yoruba-2, Vindija, Chimp) | -0.003 | -0.306 | Neither Yoruban shows increased Neanderthal ancestry over the Mbuti baseline. |
| D(Vindija, Altai, Yoruba-1, Chimp) | 0.024 | 1.095 | Neither Yoruban shows more affinity to the Vindija Neanderthal than the Altai, as would be expected if introgression came via a Eurasian vehicle. |
| D(Vindija, Altai, Yoruba-2, Chimp) | 0.034 | 1.526 | |
| D(Vindija, Altai, Mbuti-1, Chimp) | 0.002 | 0.103 | |
| D(Vindija, Altai, KhomaniSan-1, Chimp) | 0.023 | 1.008 | |
| D(Vindija, Altai, French, Chimp) | 0.054 | 2.40 | The markers in the segments retain power to identify affinity between Vindija and Europeans. |

Table S14: Summary of relevant D statistics. Putatively migrated segments were isolated in Yoruba-1 (the test Yoruban) and compared against variation in Yoruba-2 (comparable Yoruban). French, Yoruban, Han, and Mbuti individuals are from the Simons Genome Diversity Panel (sample ID `S_Sample-n.DG` in the Reich Human Origins Dataset), while Vindija and Altai are the published versions in the same dataset. Chimp (`Chimp.REF`) was used as an outgroup for all analyses.

### S3 Details of Data Analysis

#### S3.1 Inferring population size and migration rates in the Simons Genome Diversity Panel

This section describes analysis of the Simons Genome Diversity Panel with both SMCSMC and MSMC. SMCSMC version 1.0.1 was installed from the conda package manager (also found at https://github.com/luntergroup/smcsmc/releases/tag/v1.0.2), MSMC2 version 2.1.2 was installed from Github (found at https://github.com/stschiff/msmc2/releases/tag/v2.1.2) and all analyses were performed on the Oxford Biomedical Research Computation cluster.

We download prephased sequencing data from https://sharehost.hms.harvard.edu/genetics/reich_lab/sgdp/phased_data/ and mask for the strict accessibility mask from the 1000 genomes project. We additionally mask for any sites absent Chimpanzee ancestry due to a known issue with the phasing algorithm [56]. We do this masking in vcftools. We use SMCSMC to convert the sequence data from VCF to seg file format, a format very similar to MSMC format. We provide a script to convert from seg file format to MSMC file format as well. Unless otherwise noted, the names of individuals used in this paper are the first in their population (i.e. an individual named Yoruban is S_Yoruba-1 in the SGDP nomenclature, full list in S9). We select two diploid individuals from each population in Africa and infer piecewise constant population size and directional migration rates. Specifically, we use the following options for SMCSMC:

```
smc2 -c -chunks 100 -no_infer_recomb -nsam 4 -I 2 2 2 -mu 1.25e-8 -rho 3e-9 \
    -calibrate_lag 1.0 -EM {EM} -tmax 3.5 -alpha 0.0 -apf 2 -N0 14312 -Np {Np} -VB \
    ${DEMOGRAPHIC_MODEL} -P 133 133016 31*1  -arg -o ${OUTPUT} -segs ${SEGS}
```

In order, we invoke the use of a QSUB cluster with -c and split our analysis into 100 chunk. We do not infer recombination sites along with the demographic model in order to reduce runtime. Four haploid samples, two from each population, are analysed with a mutation rate of $1.25 \times 10^{-8}$, a recombination rate of $3 \times 10^{-9}$, and accumulating events for one unit of survival time along the sequence. We use a given number of epochs for parameter units, and bound the upper limits of the trees at 3.5 times the effective population size (set to 14312). We use the look-ahead likelihood to guide the resampling process for a given number of particles Np and use variational Bayes in place of the default stochastic expectation maximization algorithm. Parameters are inferred over 31 equally spaced intervals from 133 to 133016 generations in the past, and the sampled posterior ARGs are reported.

We seed the particle filter with a demographic model of population size and uniform symmetric migration rate, given by the following scrm command:

```
-ej 0.2324 2 1 -eM 0 1 -eN 0.0 6 -eN 0.0037 4.4 -eN 0.0046 3 -eN 0.0058 2 -eN 0.0073 1.4
-eN 0.0092 0.85 -eN 0.093 1.2 -eN 0.12 1.7 -eN 0.15 2.2 -eN 0.19 2.5 -eN 0.24 2.4
-eN 0.30 2.0 -eN 0.37 1.7 -eN 0.47 1.4 -eN 0.59 1.2 -eN 0.74 1.0 -eN 0.93 0.91 -eN 1.2 1.6
```

We visualise this demographic model in the POPdemog package in Figure S14 [46].

Each SMCSMC analysis gives a final output file detailing migration and coalescent events, their rates, and their opportunities which denote the total opportunity for an event to occur during a particular epoch. Output files are trimmed to only visualise the final epoch of variational Bayes inference and assessed for convergence. Times and rates are interpreted differently than scrm output. Rates are in units of $4N_0$ per generation, while times are given in generations.

We implement the above in a Snakemake pipeline. Sample size and relative cross-coalescent rates are transformed as described in the documentation using the same parameter values for mutation rate and generation time used for SMCSMC analysis. Effective population size and migration estimates for the populations analysed in the SGDP are given in Figures S5 and S2. MSMC appears to consistently find a higher African $N_e$ in the ancient past until the average estimates across populations stabilises approximately 100kya (Figure S4a). We expand on a possible reason for this effect in the main text of this article.

Migration during the last 100ky is integrated into a metric we call the inferred migration fraction (IMF). (Figure S6). This is related to the cummulative migration fraction (CMF) as introduced in MSMC-IM [56], except the quantity is integrated in a particular epoch. We use two methods to integrate migration, the first presented in the main text given by $F(t) = e^{-\int_{t=0}^{T} \rho(t)dt}$. Alternatively, consider $p$ proportion of the population are replaced every generation. Start with 0 individuals from the source $N_{source}$ population in the sink population $N_{sink}$, each generation replace $p$ proportion of the sink population with the source. We track the proportion of the population which are replaced by the source $P$.

$$
\begin{aligned}
P_0 &= 0 \\
P_1 &= pN_{sink} \\
P_2 &= pN_{sink} + p(N_{sink} - pN_{sink}) \\
&= pN_{sink} + pN_{sink}(1-p) \\
P_3 &= pN_{sink} + pN_{sink}(1-p) + p((N_{sink} - pN_{sink}) - p(N_{sink} - pN_{sink})) \\
&= pN_{sink} + pN_{sink}(1-p) + p(N_{sink}(1-p) - pN_{sink}(1-p)) \\
&= pN_{sink} + pN_{sink}(1-p) + pN_{sink}(1-p)(1-p) \\
&\dots \\
P_n &= N_{sink}p(1-p)^n
\end{aligned}
$$

In practice, both methods give essentially identical proportions for all considered questions. Inferred migration varies across language groups (Figure 1, Table S4). Afroasiatic groups show high migration from Han and French populations, with a lower proportion deriving from Papuans. Niger-Kordofanian and Nilo-Saharan groups show an intermediate magnitude, between 50 and 60 percent replacement, though also significantly (P ¡ 0.05, two-tailed paired $T$ test) closer to French and Han sources than Papuans. The Khoesan show the lowest migration, consistent with their early diversification from the remainder of African groups and the relative lack of gene-flow from Western African populations [53]. This is contrasted with the Mbuti and Biaka, Central African Hunter Gatherer populations who have historically received substantial amounts of gene flow from Western African sources. Both of these populations show the lowest migration in their language group (Table S1).

We track the overall peak of migration rate in different populations (Figure S1a,b). The most common backwards migration peak falls in the epoch between 35–45kya in the Nilo-Saharan and Niger-Kordofanian groups. Forwards migration has an earlier peak, in the epoch spanning 55–70kya. This result must be interpreted in light of the simulation results presented below.

We model the migration adjusted $N_e$ in Eurasian populations, averaged over African partners, and African populations averaged over Eurasian partners S7. The resulting curves largely represent our prior knowledge of world history, with an early divergence of Papuans consistent with the timing proposed in [34], and a second bottleneck of populations inhabiting North America such as the Karitiana and Pima. Because we do not explicitly infer population split times, and have no convenient metric like `MSMC2`, more fine-scale trends are difficult to identify. The African population size models show more discrepancy between populations, including an OoA-like bottleneck in Afroasiatic populations, and a large historical population size in hunter-gatherer groups such as proposed in [53].

#### S3.2 Validation in a physically phased subset of the Human Genome Diversity Panel (HGDP)

Phased data is not essential for demographic inference using `SMCSMC`; however, the use of phase alongside the look-ahead likelihood allows for more efficient convergence. The Human Genome Diversity Project collected 929 genomes from a diverse collection of human populations [48]. 36 of these genomes, two each from nine Eurasian and four African populations, were physically phased by use of linked-read sequencing technologies. This resource allows us to validate our inference both in an independent dataset, and evaluate the effect of phasing errors on `SMCSMC` inference.

To analyse these data, the same `Snakemake` pipeline was used with minor adjustments in wildcard constrains to account for differences in sample names. 120 chunks of the genome were run in parallel for reasons of computational efficiency, while fixed recombination rate and mutation rates were held at the same values as the SGDP analysis, and an identical demographic model was used to initiate the analysis. Three replicates of the analysis were performed to assess the impact of stochastic sampling variation on inference. We infer both effective population size and directional migration in each of these 9x4 comparisons between Eurasian and African populations (Figure S4a). The resulting inference allows us to verify and validate many observations from the SGDP.

Firstly, we calculate the timing of the migration peak and its magnitude, and find the estimates largely in line with the SGDP inference (Figure S1c,d). For instance, inferred backwards migration in the Yoruban and Biaka populations peak at 40-50kya, while the Mbuti and San show earlier migration peaks around 50-60kya (Figure S1c). The migration rate at the peak shows the same qualitative trends as the SGDP, with the peak in the Yoruban (approximately $2.5 \times 10^{-4}$) far exceeding the peaks in the Biaka, Mbuti, or San (between $0.1 - 0.175 \times 10^{-4}$) (Figure S1d). This replication in the HGDP confirms the presence of a large directional migration in the Late Middle Pleistocene, and demonstrates that statistical errors in phasing the SGDP are not large contributors to the qualitative trends observed.

We integrate migration between 40–70kya to obtain the inferred IMF for each of the comparisons in the HGDP (S4b). Differences amung the African populations mirror those in the SGDP, though the proportions are uniformly smaller (main text). However, the migration rates backwards into Africa are apparent in all comparisons, and the order of populations IMF remains the same. Differences between individual Eurasian donor populations are small, and with the exception of the Papuan, insignificant. A discussion of the Papuan comparisons appears in the subsequent section.

To compare the HGDP inference with the SGDP inference, we construct a set of the SGDP with the same donor populations as the HGDP.

#### S3.3 Comparisons between the HGDP and a subset of the SGDP

Previously, inference in the SGDP has relied on three candidate Eurasian donor populations. However, the physically phased subset of the HGDP provides a higher resolution view into global migration patterns with nine Eurasian populations represented. In order to compare effectively between the inferences made in these two datasets, we find representatives from these nine Eurasian populations in the SGDP dataset and use them as donor populations to the same four African populations (Yoruban, San, Mbuti, and Biaka), effectively recreating the analysis done in the physically phased subset of the HGDP. We select the Khomani San as a representative of the San, and only use one of the Papuan populations in the HGDP to compare (Highlands, as opposed to Sepik), creating the same 8x4 analysis table for both data sets. We infer the effective population sizes and migration rates using both `SMCSMC` and `MSMC`, with analysis details effectively identical to the original comparisons listed above (Figure S3). We average over inferences to visually compare trends between the two datasets, in the same populations and compute the inferred IMF between 40–70kya (Figure S4).

In both the HGDP and the SGDP, MSMC estimates of African population size are higher than `SMCSMC` estimates in the ancient past (80 – 300kya) (Figure S4a). By modelling directional migration, we are able to account for excess genetic diversity in the ancestral African population in both datasets. Uncertainty in the estimates increases substantially nearer to the present, as would be expected with the `SMCSMC` method.

We summarise migration from 40–70kya in the HGDP similarly to the SGDP. The total inferred migration is lower in the HGDP than in the SGDP (Figure S4b). We use this comparison setup to additionally test the differences between Papuan donors and the remainder of Eurasians. We construct a linear model predicting IMF based on an indicator variable of Papuan/not Papuan and the receptor donor population, and find that in both the SGDP and the HGDP, Papuans show approximately 2% less IMF than other donor populations (Tables S3, S2). While this difference is small, it is highly significant. However, the demographic scenario causing this difference in inferred IMF is not obvious; it is possible that the Papuan group had begun to diverge from the donating population prior to the admixture event, or alternatively that differences in archaic admixture between Eurasian and Papuan groups make up the difference in affinity.

However, the qualitative patterns in inferred directional migration rates between populations are similar in both datasets (Figure S4c). In both datasets, the highest rates are found in the Yorubans, follow by the Biaka, then the Mbuti and San. The MSMC curves are interestingly dissimilar between the different data sets, with a much steeper ascent around the period of our inferred migration in the HGDP than the SGDP.

##### S3.4 Samples used in these analyses

For a list of the identifiers used for the samples listed in this article, see Table S9.

#### S4 Statistical Analysis of Migrated Segments

We run `SMCSMC` with the `-arg` flag to report the posterior estimate of the ancestral recombination graph. We use this to isolate segments of the African genome where predicted migration events occurred between 50 and 70kya and used these segments to calculate drift statistics. The isolation procedure is implemented in `smcsmc.find_segments`, and involves sequentially reconstructing marginal trees and keeping track of which contain migration events in a particular epoch. We isolate segments from the marginal trees of all SGDP comparisons.

##### S4.1 Length Distribution of Isolated Segments

Under the Markovian model of the SMC', the length of admixed tracts $L$ is an exponential process with scale factor $2N(1-m)\left(1-e^{-T/2N}\right)$, with a proportion $m$ of the sink population being replaced with the source $T$ generations in the past and an effective population size of $N$ [7, 19]. This gives an approximate mean length $[(1-m)r(T-1)]^{-1}$ with recombination rate $r$ in units of Morgans, which is well approximated by $(rT)^{-1}(1-m)$ [27]; we use this approximation to derive expected distribution of fragment sizes. When analysing populations with `SMCSMC`, we fix the recombination rate at $3 \times 10^{-9}$ uniformly across the genome, in line with that used by tt MSMC in simulations [22, Supp. section 7]. This value is a conservative underestimate, accounting for the presence of recombination hotspots and `SMCSMC`'s inability to deconvolve recombinations in these areas, effectively underestimating the true $r$. For estimates of ancestral tract lengths, we use the more universally accepted value of $1 \times 10^{-8}$, equivalent to a one percent chance of a cross-over per megabase and per generation [9].

We take the isolated segments (Figure S11a) and compute the mean track length (Table S10). We use the approximation that the mean segment length should be approximately equal to $((1-m)r(T-1))^{-1}$ to determine that, if the migration happened in one pulse, our empirical distribution would suggest either a recent timing or a very large pulse (Figure S11b). However, we heavily caveat any interpretation of these data with the fact that they are explicitly generated under a model of a given migration proportion. The fact, therefore, that they are of a consistent length with a large migration is more evidence for the model producing internally-consistent tracts than any external validation of the results in this article.

The assumption that migration has occurred in a single wave is largely unrealistic. We used expectation maximisation to investigate if a mixture of exponential distributions explained the observed tract lengths better than a single distribution. We found that in some cases, two or three exponential distributions were better supposed by the data, however the differences in log likelihoods was negligible and the support for the different distributions was approximately inversely proportional to their number (data not shown). We found no strong support for multiple waves of migration from this analysis.

##### S4.2 Individual D statistics

Here, Yoruba-1 is used as a representative of Western African groups, and used for ascertaining putatively migrated segments. Yoruba-2 is used as a comparison individual from the same populations. In this way, we look for evidence above another individual in the same population of similarity to Eurasians.

We first use $f_3$ statistics to look for evidence of admixture between the African and various Eurasian groups. We calculate $f_3$(Yoruba-1, Eurasian group, Yoruba-2) for Papuans, French, Han Chinese, and the Vindija Neanderthal. We calculate this statistic in all available markers, and additionally for the segments isolated from the three Eurasians separately (Figure S12). These statistics show, firstly, that ascertaining in a particular group influences the shared drift with that group. This is exemplified by the non-significant shared drift with Papuans in French and Han ascertained segments. Secondly, these statistic show significant levels ($|Z| > 2$) of drift between the test individual and Eurasian populations, while also showing no increase in Neanderthal allele sharing ($f_3 = 0$). To find statistical evidence of admixture, we compute $f_3$(Yoruba-2, Eurasian, Yoruba-1) for the same Eurasian groups. We find statistical evidence for admixture in each of the groups examined, for all ascertainment schemes (Figure S13).

We use $D$ statistics to examine more nuanced scenarios. We find that the two Yorubans share more alleles than other groups in Africa (D(African group, Yoruba-1; Yoruba-2, Chimp) is significantly negative with $|Z| > 3$), but the individual of interest is closer to Out of Africa (OoA) groups such as the Han, French, and Papauns (D(OoA, Yoruba-2; Yoruba-1, Chimp) is significantly negative with $|Z| < 3$) than to its partner Yoruban (Table S5). This implies that SMCSMC has identified segments of the African Genome which are more closely related to OoA populations than to fellow Africans.

### S5 Simulation procedure

The ability of SMCSMC to recover a back-migration signal is evaluated through simulation. One gigabase of sequence was simulated in scrm, and subsequently re-inferred by SMCSMC. Migration is parameterised by three factors, magnitude, midpoint, and duration. A scenario is simulated where the midpoint is the center of a block of a given duration which has uniform migration which integrates to a given total proportion replacement over the period. We use the following demographic model for population size throughout all simulations:

The following commands can be used in either ms or SCRM to specify demographic models. The African population size is given by

```
-en 0.00000000 1 36.9124479 -en 0.00229999 1 14.8978177 -en 0.00299994 1 7.04453213
-en 0.00391291 1 3.68961222 -en 0.00510371 1 2.06587476 -en 0.00665692 1 1.21617010
-en 0.00868280 1 0.75362392 -en 0.01132521 1 0.49927968 -en 0.01477178 1 0.36258332
-en 0.01926724 1 0.29687253 -en 0.02108190 1 0.28637149 -en 0.02513079 1 0.28071694
-en 0.03277878 1 0.31028768 -en 0.03915210 1 0.36107482 -en 0.04275426 1 0.39815181
-en 0.05576555 1 0.57528787 -en 0.07273654 1 0.88701054 -en 0.09487226 1 1.36014053
-en 0.12374449 1 1.92573639 -en 0.16140334 1 2.36832894 -en 0.21052280 1 2.45284038
-en 0.27459066 1 2.16222564 -en 0.35815613 1 1.71146032 -en 0.46715286 1 1.32388966
-en 0.60932028 1 1.09778746 -en 0.79475315 1 1.04669123 -en 1.03661833 1 1.16969768
-en 1.35208972 1 1.45788656 -en 1.76356769 1 1.80077313 -en 2.30026970 1 1.89942369
```

While the European population size is given by

```
-en 0.00000000 2 1.14422216 -en 0.00229999 2 1.14422216 -en 0.00299994 2 1.14422216
-en 0.00391291 2 1.14422216 -en 0.00510371 2 1.14422216 -en 0.00665692 2 1.14422216
-en 0.00868280 2 1.14422216 -en 0.01132521 2 1.14422216 -en 0.01477178 2 1.14422216
-en 0.01926724 2 1.14422216 -en 0.02108190 2 1.14422216 -en 0.02513079 2 1.14422216
-en 0.03277878 2 1.14422216 -en 0.03915210 2 1.14422216 -en 0.04275426 2 1.14422216
-en 0.05576555 2 1.14422216 -en 0.07273654 2 1.14422216 -en 0.09487226 2 1.36014053
-en 0.12374449 2 1.92573639 -en 0.16140334 2 2.36832894 -en 0.21052280 2 2.45284038
-en 0.27459066 2 2.16222564 -en 0.35815613 2 1.71146032 -en 0.46715286 2 1.32388966
-en 0.60932028 2 1.09778746 -en 0.79475315 2 1.04669123 -en 1.03661833 2 1.16969768
-en 1.35208972 2 1.45788656 -en 1.76356769 2 1.80077313 -en 2.30026970 2 1.89942369
```

Times are in units of $4N_0g$ while population sizes are in units of $N_0$. For $g = 29, N_0 = 14312$, the demographic model is as shown in Figure S17.

The demographic model which we have assumed for both population's effective sizes has been shown to recapitulate similar inference to real data (data not shown). The migration parameter must be initiated at a given magnitude; further back in time, the particle filter is less able to identify lineage's true populations, and the inference of migration rates becomes essential uniform. Thus, we see a "drop-off" effect, where in the ancient past, the inference remains at the initiation value, and as more certainty about different histories is obtained, the migration values recapitulate real information. Thus the choice of an appropriate parameter for the initial migration rate is a crucial step in `SMCSMC` analysis, and here we chose to arrive at this value through simulation.

We simulate back-migration scenarios of varying total migration proportions from 0 (no migration) up to 60% population replacement.For each simulation, we initiate the particle filter at either 0, 1, or 5 $4N_0$ proportion replaced per generation (which are the units used internally by `scrm` and `ms` for simulation). `SMCSMC` is then used to infer effective population size and migration histories in five iterations with 5000 particles. As a cautionary note, these simulations are almost certainly not fully converged, and are used as an indication of power. Their power, theoretically, approaches 1, as particle filters asymptotically exactly approach the true posterior distribution. However, these low resolution attempts are indicative of a "quick" overview of the abilities of the algorithm. With 600 cores available, each of the cases (forward, backward, or bidirectional) was able to run in approximately 20 hours.

Generally, beginning with a higher migration rate seems to recover a higher proportion of the simulated migration. However, as in the case of a 60% replacement simulated 40kya, beginning with 5 $4N_0$ rather than 1 $4N_0$ recovers similar proportions of backwards migration (0.502 vs 0.52) yet the higher migration rate finds 0.301 Eurasian migration rather than 0.195. The higher initial migration rates thus slightly reduce power (though, not in all cases, and for fully converged solutions, we would expect both proportions to be similar up to noise) while additionally finding an increased migration in the opposite direction. Beginning with a zero rate leads to highly unstable estimates of the migration rate and effective population size, and we exclude it from our analysis.

We select a more comprehensive set of initiation parameters and particle values and use them to analyze a Yoruban and French individual from SGDP (Fig S16). The effect of the initial migration rate seems relatively consistent for low values (0.5 - 2.0),while an increasingly small migration peak is seen for higher initial magnitudes 4.0 - 10.0. Again, beginning with an initial rate of zero tends to lead to highly unstable estimates of effective population size and migration rates. For the remainder of the analyses in this article, we choose to use an initial rate of 1.0.
